# Plasmodium falciparum subverts neutrophil function via host miR-451a loaded extracellular vesicles driving bacterial superinfection susceptibility

**DOI:** 10.1101/2025.06.02.657463

**Authors:** Kehinde Adebayo Babatunde, Mtakai Ngara, Paola Martinez Murillo, Bibin Subramanian, Alex Hopke, Simone Lucchesi, Maria Novedrati, Giorgio Montesi, Francesco Santoro, Michael Hagemann-Jensen, Rickard Sandberg, Luis Filgueira, Michael Walch, Daniel Irimia, Pierre-Yves Mantel

**Affiliations:** The Faculty of Science and Medicine, Department of Oncology, Microbiology and Immunology, University of Fribourg, 1700, Fribourg, Switzerland; Christine Kühne-Center for Allergy Research and Education (CK-CARE), Davos, Switzerland; Department of Pathology & Laboratory Medicine, University of Wisconsin, Madison, WI 53705, USA; Department of Microbiology, Tumor and Cell Biology, Karolinska Institutet, Sweden; Department of Surgery, BioMEMS Resource Center, Massachusetts General Hospital, Harvard Medical School, Boston, MA, USA; Department of Cell and Molecular Biology (CMB), Karolinska Institutet, Sweden; Department of Medical Biotechnology, University of Siena, 53100 Siena, Italy

**Keywords:** Malaria, *Plasmodium falciparum*, extracellular vesicles, neutrophils, microRNAs

## Abstract

Malaria caused by Plasmodium falciparum (*Pf*) compromise innate immunity, yet the underlying mechanisms remain elusive. The immune dysregulation caused by the parasite may lead to bacterial superinfections and increase mortality. We reveal that *Pf* exploits extracellular vesicles (EVs) secreted by infected red blood cells (iRBC-EVs) to deliver host-derived miR451a to human neutrophils, impairing their antimicrobial defences. Neutrophil phagocytosis of iRBC-EVs suppresses reactive oxygen species (ROS) production and compromised microbicidal activity against *Salmonella typhimurium*. Microfluidic assays show that miR451a transfer significantly disrupts neutrophil chemotaxis and swarming upon microbial challenge. Transcriptomic profiling indicates that EVs and miR451a reprogram neutrophil gene expression, notably upregulating ferroptosis-related genes, suggesting a role in further impairing immune responses. We have uncovered a novel mechanism of iRBC-EVs-induced neutrophil immune suppression and provide insights into increased susceptibility to bacterial superinfections in malaria. These findings have implications for therapeutic strategies aimed at mitigating bacterial superinfections and sepsis in malaria-endemic regions.

## Introduction

Malaria remains a significant global health challenge, with nearly half of the world’s population at risk of infection (Snow et al., 2005; Venkatesan, 2024). In 2022, there were an estimated 249 million cases of malaria, leading to 608,000 deaths worldwide. Alarmingly, the majority of these deaths was in children younger than the age of 5 (Organization., 2022). Beyond causing direct mortality, malaria indirectly contributes to deaths by compromising the host immune response and heightening vulnerability to secondary infections from bacteria or fungi. Indeed, infection with *Plasmodium falciparum*, the most virulent malaria parasite species, has been linked to an elevated risk of invasive bacterial infections, which are associated with high mortality rates among children suffering from severe malaria in sub-Saharan Africa (Were et al., 2011). While the correlation between malaria and fungal infections is poorly documented, there is an increasing number of clinical reports on fungi associated malaria infection (Dabritz et al., 2011; Eckerle et al., 2009; Hocqueloux et al., 2000; Ruhnke et al., 2000; Wilson et al., 2000). The underlying mechanism for the increased risk of co-infection is not fully understood.

Neutrophils are important in pathogen defense through processes such as inflammation, NETs formation, ROS production and swarming behavior around invading pathogens (Hopke et al., 2020). However, malaria infection has been linked to defective neutrophil responses. For instance, malaria patients often exhibit significantly reduced neutrophil counts (Kotepui et al., 2014), and neutrophils in these patients have been reported to exhibit reduced oxidative burst, reduced pathogen phagocytosis (Cunnington et al., 2011; Cunnington et al., 2012) and impaired chemotaxis (Leoratti et al., 2012). This may partly explain the increased susceptibility to bacterial superinfection observed during malaria. While the exact cause of neutrophil impairment during malaria is ill defined, several hypotheses have been formulated such as the increase of free heme in circulating blood (Cunnington et al., 2011) and release of digestive vacuoles from the iRBCs (Dasari et al., 2012; Dasari et al., 2011).

During blood stage, the parasite releases membranous material in the form of extracellular vesicles (EVs), which are small vesicles ranging from 30 to 150 nm (exosomes) and 150 nm to 1–2 μm (microvesicles) in diameter. In severe malaria cases, the number of EVs derived from malaria-infected red blood cells (iRBCs) in patient serum is significantly elevated (Nantakomol et al., 2011). EVs are known to shuttle biological materials such as cytosolic proteins, lipids, DNA and RNA. Beside their role in immune regulation, EVs have been reported to aid cell-to-cell communication between *P. falciparum* parasites to synchronize the transmission stage (Regev-Rudzki et al., 2013), and between *P. falciparum* and host (Mantel et al., 2013). Here, we hypothesized that circulatory iRBC-EVs could play an important role in the observed neutrophil dysfunction during malaria. Our data show that iRBC-EVs transfer the host miRNA, miR451a to neutrophils resulting in several functional impairments. These results provide new insights into the molecular basis of neutrophils dysfunction in malaria and highlight the role of EVs as key mediator of host-parasite interactions.

## MATERIALS AND METHODS

### Fungi

*Candida albicans* SC5314 constitutively expressing a far-red fluorescent protein was cultured overnight at 30°C with shaking in YPD broth media (Hopke et al., 2016).

### Plasmodium falciparum in Vitro Culture

The 3D7 *P. falciparum* strain was kept in fresh type O^+^ human erythrocytes suspended at 2% hematocrit in HEPES-buffered RPMI 1640 (Sigma-Aldrich), containing 10% (w/v) heat inactivated human serum, 0.5 mL gentamicin (Sigma-Aldrich), 2.01 g sodium bicarbonate, and 0.05 g hypoxanthine at pH 6.74. Prior to culture, the complete medium was depleted of EVs and debris by ultracentrifugation at 100,000× *g* for 1 h before the culture. The parasite cultures were maintained in a controlled environment at 37 °C in a gassed chamber at 5% CO_2_ and 1% O_2_.

### Purification of EVs Derived from Malaria-Infected Red Blood Cells (iRBC-EVs)

EVs from iRBCs were isolated from cell culture supernatants as previously described (Mantel et al., 2013). First, 15 ml of cell culture supernatants of *P. falciparum*-infected RBCs were collected. Cells and cellular debris were removed from the supernatant by centrifugation at 600× *g*, 1600× *g*, 3600× *g*, and, finally, 10,000× *g* for 15 min. To further concentrate the EVs, the supernatant was filtered through a Vivacell 100 filter (100 kDa molecular weight cut off; Sartorius). Then, the concentrated supernatant was pelleted at 100,000× *g*, the pellet resuspended in PBS and layered on top of a 60% sucrose cushion and spun at 100,000× *g* for 16 h. The interphase was collected and washed with PBS twice at 100,000× *g* for 1 h to yield EVs.

### Neutrophil isolation from whole blood

Human blood samples from healthy donors (aged 18 years and older) were purchased from Research Blood Components, LLC. Primary neutrophils were isolated within 1h after draw using the human neutrophil direct isolation kit (STEMcell Technologies) following the manufacturer’s instructions. The purity was shown to be higher than 96% by flow cytometry. For microfluidic experiments, primary neutrophils were stained with Hoechst 33342 trihydrochloride dye (Thermo Fisher Scientific). Stained neutrophils were then suspended in RPMI 1640 media containing 20% FBS (Thermo Fisher Scientific) at a concentration of 2 × 10^7^ cells/mL in the chemotaxis device or 2.5 x 10^6^ cells/mL in the swarming assay.

### Confocal analysis for internalized EVs

Neutrophils (2 x 10^6^ cells/mL) were seeded onto coverslips treated with 0.01% poly-L-lysin for 30 min at room temperature (RT) in a 24-well tissue culture plates, incubated at 37°C in a humidified 5% CO_2_ atmosphere. Cells were then washed three times before inhibitor pre-treatment for 30 min, before adding 100μg of PKH67 labeled EVs to the neutrophils and incubated for 2 hours. The following inhibitors were used: Dynasore (80 μM), colchicine (31 nM), latrunculin A (5 μM) and nocodazole (20 μM). Next, the excess and unbound EVs were removed by 3 washes with PBS and the cells were fixed with 3% paraformaldehyde in PBS for 15 minutes at RT. After fixation, cells were permeabilized for 5 minutes in PBS containing 0.1% Triton X-100, washed with PBS, and stained with phalloidin dye (Invitrogen) for 20 min at RT then washed off gently 3 times with PBS afterward to remove excess phalloidin dye. The DNA was counterstained for 5 minutes with Hoechst 33342. Cover-slips were mounted on a glass slide with mounting media. Images were captured using a Leica TCS SP5 confocal microscope with a 63X oil immersion objective. Images were collected, and confocal *z*-sections at 1024 X 1024 magnification were acquired at 0.3-mm intervals. Images were further analyzed and processed using Imaris software, Image J and affinity designer software.

### Imaging flow cytometry

Imaging flow cytometry was performed on an ImageStream^X^ Mark II operated by INSPIRE software (Amnis Corporation). PKH-67 fluorescence was recorded using excitation with a 488nm laser at 50 mW intensity and emission collected with a 480nm-560nm filter (CH2), and DRAQ5 fluorescence using excitation with a 642nm laser at 30 mW intensity and a 660nm-740nm filter (CH5). Brightfield images were collected in CH 1. A total of 10’000 events were collected for each sample. Single stained controls were also collected (PKH-67 EVs only and DRAQ5 only stained cells) at the same settings, to develop a compensation matrix for removing spectral overlap of the dyes from each of the channels. Data were analyzed using IDEAS Application v6.0 software (Amnis Corporation). A compensation matrix was created using single color controls and applied to all files. Image-based gating was performed as follows: Isolated primary human neutrophils were incubated with PKH67-labeled EVs for 1h. After identification of focused cells D(i) and single cell events D(ii), a population of cells with high EVs intensity were identified and excluded from further analysis D(iii) and D(iv). The remainder of the cells were named low EVs intensity D(iii) and D(v). Orange = Low EVs intensity population, Yellow = High EVs intensity population. The internalization feature was used to identify neutrophils containing no PKH67-labeled EVs or containing PKH67-labeled EVs from the Low EVs intensity population D(vi). The Spot Count feature was used to identify neutrophils containing no PKH67-labeled EVs and neutrophils containing 1 to 2 PKH67-labeled EVs from the Low EVs intensity population D (vii) and (viii).

### Killing experiment in neutrophils and HL-60

*Ex vivo* killing of bacteria by blood neutrophils was assessed in a gentamicin protection assay and quantified by bacterial culture. *Salmonella typhimurium strain SL1344* or *Staphylococcus aureus* were cultured overnight in 10 mL LB at 37 °C with shaking at 175 rpm. The next day, the bacteria were diluted 1:20 and let grown to the exponential phase (OD=600). The bacteria were opsonized for 10 min in human serum. Cells were resuspended at a density of 1×10^6^/mL in RPMI without antibiotics and seeded at 5×10^5^ cells per well in a 24-well. Plates were incubated at 37°C and 5% CO_2_ for 20 minutes prior to the addition of *S. typhimurium* or *S. aureus* at a multiplicity of infection 10:1 (MOI, bacterium: eukaryotic cell ratio). After the incubation time, the medium was removed and replaced with medium containing gentamicin (10 μg/mL in RPMI) for 30 minutes to kill extracellular bacteria. The cells were washed and lysed in 500 μl of ice-cold distilled water containing 1% Triton for 1 hour for lysis of neutrophils and 10-fold dilutions were plated on LB agar, incubated for 18 h at 37°C and viable bacteria colonies were quantified using Image J. Growth experiments were repeated at least three times.

### Generation of HL60^miR451a^ and HL60^scramble^ lines

For the construction of the miR451a lentiviral vector, fragments containing the human miR451a genomic clusters were amplified by PCR from human genomic DNA and directionally cloned into the *AgeI* and *EcoRI* sites of the pLKO-1 vector (Addgene#8453, Cambridge, MA) using the primers: forward 5′-AAAAACCGGTCTAGTCCGGGCACCCCCAG −3′ and reverse 5′-AAAAGAATTCCCTACCCCCAATCCCACGC −3′. Lentiviral particles were produced by transfecting HEK 293F cells using the LV-MAX Lentiviral Production System with packaging plasmid psPAX2 and envelope plasmid pMD2.G (both Addgene#12260, #12259) and harvesting supernatants after 48 h, as described by manufacturer’s instructions. Viral supernatant was harvested 48 h post transfection, filtered (0.45 μm pore size), and transduced into HL-60 cells in the presence of 8 μg/mL of polybrene (Sigma-Aldrich). After 24 h, cells were maintained in medium containing 0.6 μg ml^−1^ puromycin (Sigma-Aldrich) for selection. As a negative control we used the plasmid pLKO.1 GFP shRNA Addgene#30323) with the target sequence 5’GCAAGCTGACCCTGAAGTTCAT3’.

### RNA isolation and sequencing

Total RNA was extracted from 5×10^6^ neutrophils or 1×10^6^ HL-60 cells using the RNeasy Mini Kit including a DNase digest according to the manufacturer’s instructions (QIAGEN). The quality and quantity of the extracted RNA were assessed using a NanoDrop spectrophotometer and a Qubit fluorometer (Thermo Fisher Scientific). RNA integrity was further evaluated using the Agilent 2100 Bioanalyzer (Agilent Technologies), ensuring that all samples had an RNA integrity number (RIN) >7.0 before proceeding to library preparation.

For library preparation, the mRNA present in the total RNA sample was isolated with magnetic beads of oligos d(T). Subsequently, mRNA was randomly fragmented, and cDNA synthesis was performed using random hexamers and reverse transcriptase enzyme. The library was ready after end repair, A-tailing, adapter ligation, size selection, amplification, and purification. The library preparation was performed using Novogene’s property RNA Library Prep Kit (PT042) based on NEB Next® Ultra™ RNA Library Prep Kit and NEB Next®.

The library was checked with Qubit 2.0 and real-time PCR for quantification and bioanalyzer Agilent 2100 for size distribution detection. Quantified libraries were pooled and sequenced on Illumina NovaSeq X Plus. The NovaSeq X Plus sequencing system was used to sequence the libraries in a paired end 150bp (PE150) with 6 GB raw of data output per sample.

Single-end cDNA libraries were prepared for each sample using TruSeq RNA Sample Preparation Kit and sequenced on Illumina’s HiSeq 2000 platform. The data for RNA-Seq were uploaded to GEO (GEO accession number: GSE296044).

### Microfluidic device fabrication

To prepare the glass swarming assay, ultraclean glass slides (ThermoFisher UltraClean Microarray Slides, Fisher Scientific) were micro-patterned with a solution containing poly-L-lysin and FITC-ZETAG (1.6mg/mL) using a Polypico micro-dispensing machine (PolyPico Technologies). Live *Candida albicans* were used as targets for neutrophils swarms. *Candida albicans* were cultured overnight and suspended in sterile water prior to micro-patterning. *C. albicans* was seeded onto each glass slide and was allowed to adhere for 5-10 mins on a rocker. Excess *C. albicans* was then washed twice with PBS. Prior to experiment, glass slides were placed in a commercially available open well chamber (Hopke et al., 2020).

The microfluidic devices for NETs assay were fabricated using standard microfabrication technologies. One layer (35 µm thick) of SU8 photoresist (Microchem) was patterned on a silicon wafer using photolithography masks, following standard processing recommendations from the manufacturer. The wafer with patterned photoresist was used as a mold to produce pieces of polydimethylsiloxane (PDMS, Fisher Scientific), which were subsequently bonded irreversibly to standard glass slides (1 × 3 inches, Fisher).

Prior to use, microfluidic devices were primed with phosphate buffered saline (1X PBS pH 7.4, Life technologies). Blood samples (20 µL), within 1 h after collection, were diluted in 480 µL of PBS and 0.50 µL of 1 mM Sytox-Green (Thermo Fisher Scientific). The diluted blood was loaded into the large inlet reservoir while the outlet of the device is connected to a 1 mL syringe via a tube mounted on an electric pump. A volume of 100 µL diluted blood was pipetted into the outlet and pumping was done at a flow rate of 10 µL/min. The final volume processed by the device is 50 µL.

The microfluidic device used for chemotaxis assays consists of a large egg-shaped chamber with a single entrance channel connected to the inner central reservoir (Figure 3A) (Ellett et al., 2019). A chemoattractant gradient is established in the device from the large egg-shaped chamber through the connecting channel to the inner chemoattractant chambers (supplemental video 1). This device enables us to monitor and track chemotaxis in primary neutrophil towards chemoattractant gradient. To estimate number of chemotactic neutrophils using the egg-shaped device, we calculated the percentage of cells migrating into the inner chemoattractant chamber.

### Visualization and quantification of NET formation in whole blood

The quantification of NETosis in whole blood using the NET microfluidic device was carried out by measuring the Sytox-green fluorescence area on the device. Sytox-green fluorescence area was recorded by taking stitched images of the entire device using the FITC channel on a time lapse microscope at RT. The areas of chromatin trapped in the device was quantified after automatic thresholding using Image J (triangle threshold filtration, U.S. National Institutes of Health, Bethesda, MD, USA) by using the analyze particle plugin in Image J.

### Visualization of NET formation in isolated primary neutrophils

The visualization of NETosis was carried out as previously described (Brinkmann et al., 2010). Briefly, isolated neutrophils were fixed for 30 min at RT in 3% paraformaldehyde, permeabilized with 0.5% Triton X-100, and blocked for 30 min in blocking buffer. The samples were then incubated for 60 min with the primary antibody as follow: anti-human histone H3 mouse monoclonal antibody (diluted 1:100) and anti-human Cit-H3 rabbit polyclonal antibody (1:100) (ab5103; Abcam, Cambridge, UK). After washing in PBS, each primary antibody was visualized using secondary antibodies coupled to 1:500 Alexa Fluor 546 goat anti-mouse IgG (Thermo Fisher Scientific) and 1:500 Alexa Fluor 488 goat anti-rabbit IgG (Thermo Fisher Scientific). After incubation for 60 minutes with the secondary antibodies, the specimens were washed with PBS, and the DNA stained with 49,6-diamidino-2-phenylindole (Thermo Fisher Scientific) in PBS for 5 min. All procedures were performed at RT. The specimens were analyzed using a confocal laser-scanning microscope and processed using ImageJ/Fiji software.

### Analysis of primary neutrophil migration

We used Track-mate module in Fiji ImageJ (ImageJ, NIH) to track and analyze primary neutrophils migrating from the large egg-shaped chamber through the entrance channel connected to the chemo-attractant inner central reservoir. Primary neutrophils with fluorescently stained iRBC-EVs were identified by using the track mate plugin in Image-J before and after migration. Percentage of migration was calculated as thus: (N_TIC_)**/**(N_TC_)*100. Where N_TIC_ is total number of migrated cells in the chemo-attractant inner central reservoir after migration and N_TC_ is total number of cells in the device before migration.

### Swarming experiment

All imaging experiments were conducted using a fully automated Nikon TiE microscope. Time-lapse imaging was conducted using a ×10 Plan Fluor Ph1 DLL (NA = 0.3) lens. Swarming targets to be observed during time lapse were selected and saved using the multipoint function in NIS elements prior to loading of neutrophils. 200-300 hundred thousand neutrophils were added to each well unless otherwise noted. All selected points were optimized using the Nikon Perfect Focus system before starting the experiment. Changes in swarm size over time were estimated using track mate plugin in Image J. The cell-occupied area was measured from the Hoechst labeled primary neutrophil using filter and suitable threshold on image J.

### Single Cell RNA-seq

A total of 192 single cells were sorted in 96 well plates containing lysis buffer using FACS Aria and libraries generated according to Smart-seq2 protocol (Picelli et al., 2013). These included 48 single cells from each of the four experimental groups namely: HL60^miR451a^Bact, HL60^mir451a^, HL60^Scramble^bact and HL60^Scramble^. Libraries were sequenced on Illumina’s NextSeq 550 platform and the raw reads mapped towards the human reference genome (hg19) using STAR mapper (Dobin et al., 2013). Gene expression was quantified based on uniquely mapped reads as guided by Ensembl annotations. To filter low quality cells different metrics were assessed including, number of detected genes (at least 50 detected genes per cell), ratio of exon-intron mapped reads (greater than 50%), fraction of mitochondrial gene (less than 20%), fraction of reads attributable to the highly expressed genes among others. Only cells that passed quality control analysis and without detectable batch effects were used for downstream analysis. Here individual cell’s transcriptome was normalized to counts per 100K and log transformed. Using variable genes, individual cells were transformed into low dimensional space using Principal Component Analysis (PCA) analysis and projected onto UMAP/tSNE subspace. Clusters were detected using Louvain algorithm for each subpopulation. The quality control analysis, subpopulation discovery and exploration were carried out using the methods as implemented in scanpy (version 1.8.1) (Wolf et al., 2018). We used nonparametric Wilcoxon rank-sum test for differential gene expression analysis between subpopulations.

### Reactive oxygen species (ROS) assay

ROS production was assessed by measuring extracellular hydrogen peroxide release using Amplex UltraRed (Thermo Fisher) in the presence of 10 U/mL horseradish peroxidase. Neutrophils were resuspended in HBSS at a concentration of 1 × 10⁶ cells/mL. In a black 96-well flat-bottom microplate (Costar), 100 µL of cell suspension was added per well. Amplex UltraRed was used at a final concentration of 50 µM, and HRP was added at 0.1 U/mL. The cells were pre-treated or not with EVs for 1 hour. For stimulation, PMA (100 nM final concentration) was added to the wells. Control wells received vehicle alone. The plate was incubated at 37°C in the dark, and fluorescence was measured at 530 nm excitation and 590 nm emission using a microplate reader (Synergy, H1, Biokek).

### RNA Fluorescence *in situ* hybridization

We performed RNA fluorescent *in situ* hybridization staining essentially as described (Lu and Tsourkas, 2011). Briefly, we used 10 nM of DIG-labelled miRNA LNA probes (Exiqon). A scrambled miRNA LNA probe was used as a negative control. After a series of post-hybridization washes, the LNA signal was amplified using the Tyramide Signal Amplification PLUS Fluorescein Kit (Perkin-Elmer, Waltham MA) according to the manufacturer’s instructions.

### Statistical analysis

Statistical significance of the differences between multiple groups were tested using two-way Analysis of Variance (ANOVA) in GraphPad Prism (GraphPad Software, version 8.3.0). Within ANOVA, significance between two sets of data was further analyzed using two-tailed **t**-tests. Differential gene expression analysis was performed with the R package edgeR (v4.0.16) (Robinson et al., 2010). GO term enrichment analysis was performed using the ClusterProfiler R package (v3.14.3) (Yu et al., 2012). Since most genes did not show a normal distribution, differences in gene expression were assessed with the Wilcoxon test, corrected for multiple testing with the Benjamini-Hochberg method, on vst (variance stabilizing transformation) normalized data from edgeR.

## RESULTS

### Neutrophils internalize iRBC derived EVs

iRBC-EVs are a heterogenous group of cell-derived membranous structures that have a size ranging from 50 nm to 1 μm in diameter (Babatunde et al., 2020). We isolated EVs from *in vitro* cultured *P. falciparum* parasites. In line with previous studies, transmission electron microscopy demonstrated round nano-sized structures surrounded by a lipid bilayer (Figure 1A). To further confirm the size distribution of EVs, we used the nano tracking instrument that demonstrates that EVs have a size between 50 – 400 nm (Figure 1B). Next, to investigate the capacity of neutrophils to internalize iRBC-EVs, we cultured isolated primary human blood neutrophils in the presence of serum opsonized EVs for 1 hour and performed confocal fluorescence microscopy. EVs are taken up by neutrophils and are not only sticking to the cell membrane (Figure 1C). The efficient uptake and interaction were further confirmed and quantified by imaging flow cytometry (Figure 1D). We distinguished two populations of cells with High EV intensity and Low EV intensity. This might be caused by the amount of EVs phagocytosed (Figure 1E) or the heterogeneity of the EVs. To gain insight into the mechanisms of EV uptake, we pre-treated neutrophils with a series of known endocytosis inhibitors (Figure 1F). The endocytosis inhibitors tested included actin filament inhibitor: Latrunculin A, the dynamin inhibitor: dynasore, the microtubule inhibitors nocodazole and colchicine (Lamaze et al., 1997; Vasquez et al., 1997). All tested endocytosis inhibitors beside nocodazole demonstrated significant inhibitory effects on iRBC-EV uptake. Notably, EV uptake was most strongly suppressed by inhibitors targeting actin polymerization and dynamin activity.

**Figure 1:**
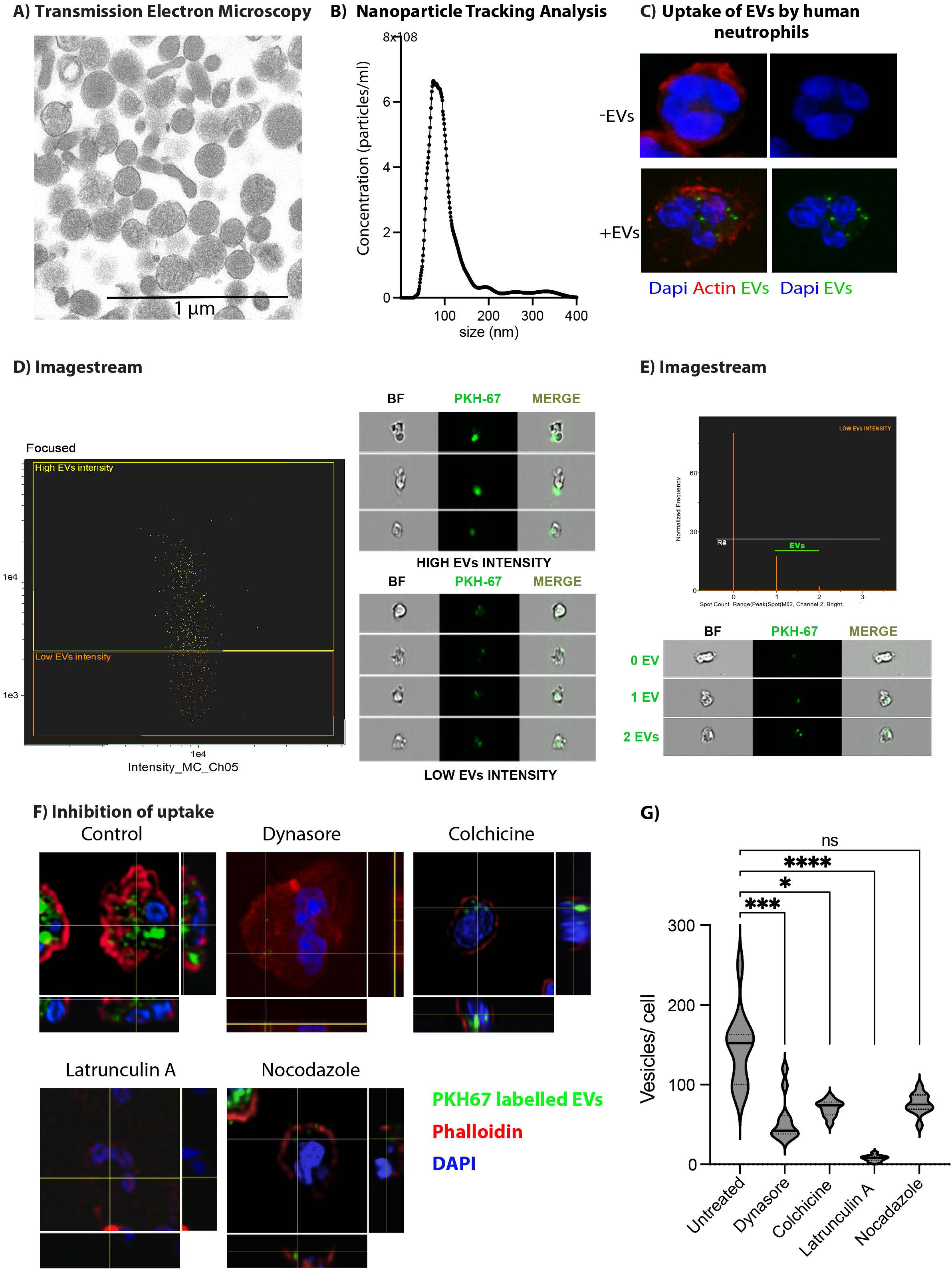
iRBCs-EVs are internalized by primary neutrophils. **A)** Transmission Electron Microscopy image of iRBC-EV. The bar represents a size of 1 μm, **B)** Size distribution of iRBC EVs as determined by Nanoparticle Tracking Analysis showing the size distribution of iRBC-EV. **C)** Confocal microscopy of EV uptake by neutrophils. Isolated neutrophils were incubated with 100μg of PKH-67 fluorescently labelled EVs for 1h at 37°C. Neutrophils were stained for actin (Phalloidin, red) and nuclei (Hoechst-blue). **D)** Representative image from Amnis ImageStream System analysis based on the green fluorescent intensity, we divided into high EV intensity (Yellow colored-top image) and low EV intensity (Orange colored-bottom image) to exclude debris. Representative images from Amnis ImageStream System analysis showing neutrophils (brightfield) interacting with **D)** fluorescently labeled iRBC-EVs (green, PKH67) (High EV intensity) and **D)** fluorescently labeled iRBC-EVs (green, PKH67) (low EV intensity). **E)** Representative image from Amnis ImageStream System analysis showing quantification of low EV intensity internalized by neutrophils. **E)** Representative image from Amnis ImageStream System analysis showing neutrophils (brightfield) with 0, 1 or 2 internalized fluorescently labeled iRBC-EVs (green, PKH67) (low EV intensity). Representative results from at least three experiments are shown. **F)** Effect of endocytosis and cytoskeletal reorganization inhibitors on EV uptake. Cells were pretreated for 30 min, with the inhibitors before adding the EVs for 2h. i) Control, ii) Dynasore (80 μM), iii) Colchicine (31 nM), iv) Latrunculin A (5 μM) and iii) Nocodazole (20 μM). One representative experiment of 3 independent experiments is shown. **G)** effect of inhibitors measured as number of EVs per cell (median; *n*=3 experiments). Comparisons versus the untreated control were performed using a one-way ANOVA with the Kruskla-Wallis test. Significance levels are indicated as \**P*<0.05, \*\**P*<0.01 and *** *P*<0.001.

### iRBC-EVs suppress immune response of human neutrophils

Neutrophils are known to produce reactive oxidative species (ROS) to kill a diverse array of pathogens. To assess how internalized EVs affect ROS production, we incubated freshly isolated neutrophils with serum-opsonized EVs for 1 hour, followed by stimulation with PMA. Our results show a significant difference in extracellular ROS production between neutrophils stimulated with PMA alone and those pretreated with EVs prior to PMA stimulation. Specifically, pretreatment with EVs substantially reduced and delayed ROS generation compared to stimulation with PMA alone (Figure 2A). To evaluate the bactericidal capacity of EV-treated neutrophils, we incubated freshly isolated neutrophils with EVs for 1 hour at 37°C. Subsequently, these neutrophils were challenged with *Salmonella typhimurium* to assess the direct bactericidal activity of EV-treated neutrophils using a colony forming unit (CFU) assay.

**Figure 2:**
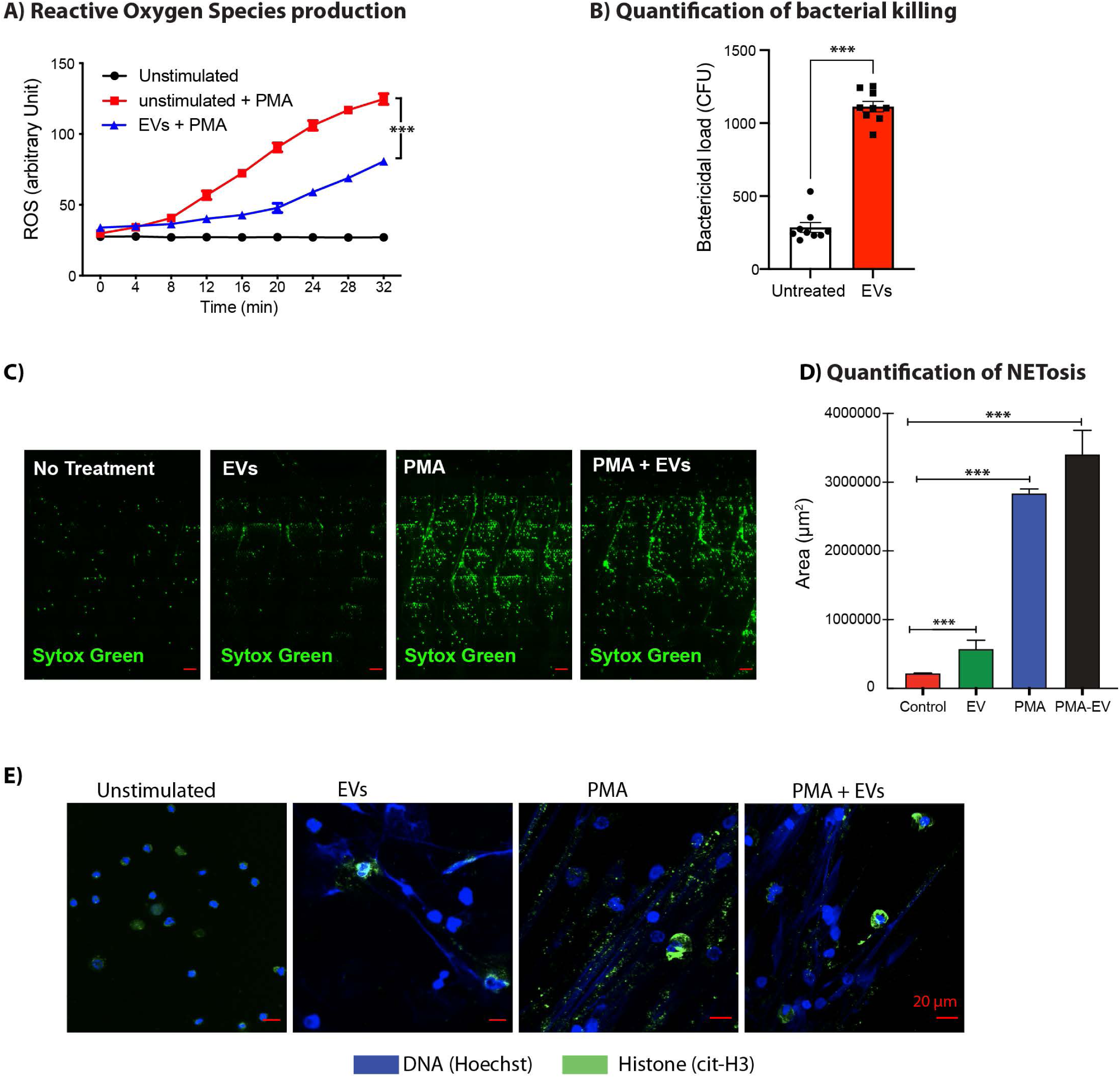
iRBC-EVs inhibit ROS production and killing abilities in primary neutrophils. **A)** ROS-production by unstimulated, PMA and iRBC EVs + PMA treated primary neutrophils stimulated with PMA. One representative experiment of 3 independent experiments performed in triplicates is shown. **B)** To determine the anti-bacterial activity, isolated neutrophils were pre-exposed to EVs or not and then incubated with *S. typhimurium* before assessing bacterial load by colony forming unit (CFU) assay. Data are expressed as means ± SEM of 3 independent experiments performed in triplicates. Statistical significance was determined using the Mann-Whitney unpaired test, with *P < 0.05, **P < 0.01, and ***P < 0.001. **C)** Representative images used for NET quantifications, showing staining for trapped extracellular DNA (green) in response to PMA (100nM), iRBC-EVs(100μg) and PMA/iRBC-EVs in whole blood from healthy donors (*n* = 4). Scale bars, 50 μm **D**) Quantification of NETosis in whole blood from healthy donors (*n* = 4) in response to PMA (100nM), iRBC-EVs(100μg) and PMA/iRBC-EVs. Asterisks indicate significance: \*\*\**P* < 0.001 by two-way analysis of variance (ANOVA). **E)** Representative images showing staining for DNA (blue) and histone (green) in response to PMA (100nM), iRBC-EVs (100 μg) and PMA/iRBC-EVs in PMNs isolated from healthy donors (*n* = 4) Scale bars, 20 μm.

Neutrophils pre-treated with EVs derived from iRBC-EVs were exposed to *S. typhimurium* at a multiplicity of infection (MOI) of 1:10 for 30 minutes. This assay allowed us to quantify the number of viable bacterial colonies following neutrophil-mediated killing, providing a measure of the neutrophil’s bactericidal efficiency. Our CFU assay revealed about 3-fold increase in viable bacteria colonies in iRBC-EV treated neutrophil compared to untreated, suggesting the reduced capacity of iRBC-EV treated neutrophils to kill the ingested bacteria (Figure 2B).

To test whether malaria EVs induce NETs formation, we used a microfluidic device. We co-incubated neutrophils from healthy adult donors with EVs derived from iRBC for an hour. PMA was used as positive control (Figure 2C). We found that PMA, EVs and PMA-EVs robustly induced NETs in whole blood. However, the mean NETs area of the trapped extracellular DNA was 2 folds larger for EVs compared to control, at 500’000 μm^2^ vs 240’000 μm^2^, respectively. The mean NETs were larger for PMA-EVs compared to PMA alone at 3.5 x 10^6^ μm^2^ vs 2.8 x 10^6^ μm^2^, respectively (Figure 2D). To further, confirm NETs stimulation by EVs, we stimulated purified neutrophils with serum opsonized iRBC-EVs for an hour and stained for extracellular DNA and citrullinated histone prior to assessment by confocal microscope. We found that iRBC-EV could also directly induce NETs (Figure 2E).

### iRBC-EVs alter chemotaxis and swarming in isolated human primary neutrophils *ex vivo*

A previous study has shown that neutrophils isolated from individuals infected with *P. vivax* malaria exhibit reduced chemotactic activity, although the underlying mechanism for the impairment is unclear (Leoratti et al., 2012). To assess whether iRBC-EVs impair neutrophil chemotaxis, we treated isolated neutrophils with serum-opsonized iRBC-EVs for 1 hour and quantified migration toward 100 nM formyl-methionyl-leucyl-phenylalanine (fMLP) using an egg-shaped microfluidic platform. This system enables real-time, single-cell resolution tracking of chemotaxis by directly measuring neutrophil migration toward chemokine gradients, a critical functional readout of chemotactic activity (Figure 3A, supplemental video 1). We found that both iRBC-EV treated and untreated neutrophils in the outer chamber migrate directionally into the central inner micro-chamber in response to the fMLP gradient. However, the mean percentage of migration was twice as much in untreated neutrophils compared to iRBC-EV treated neutrophils, at 30% vs 15%, respectively suggesting impairment of neutrophil chemotactic response following treatment with iRBC-EV (Figure 3B).

**Figure 3:**
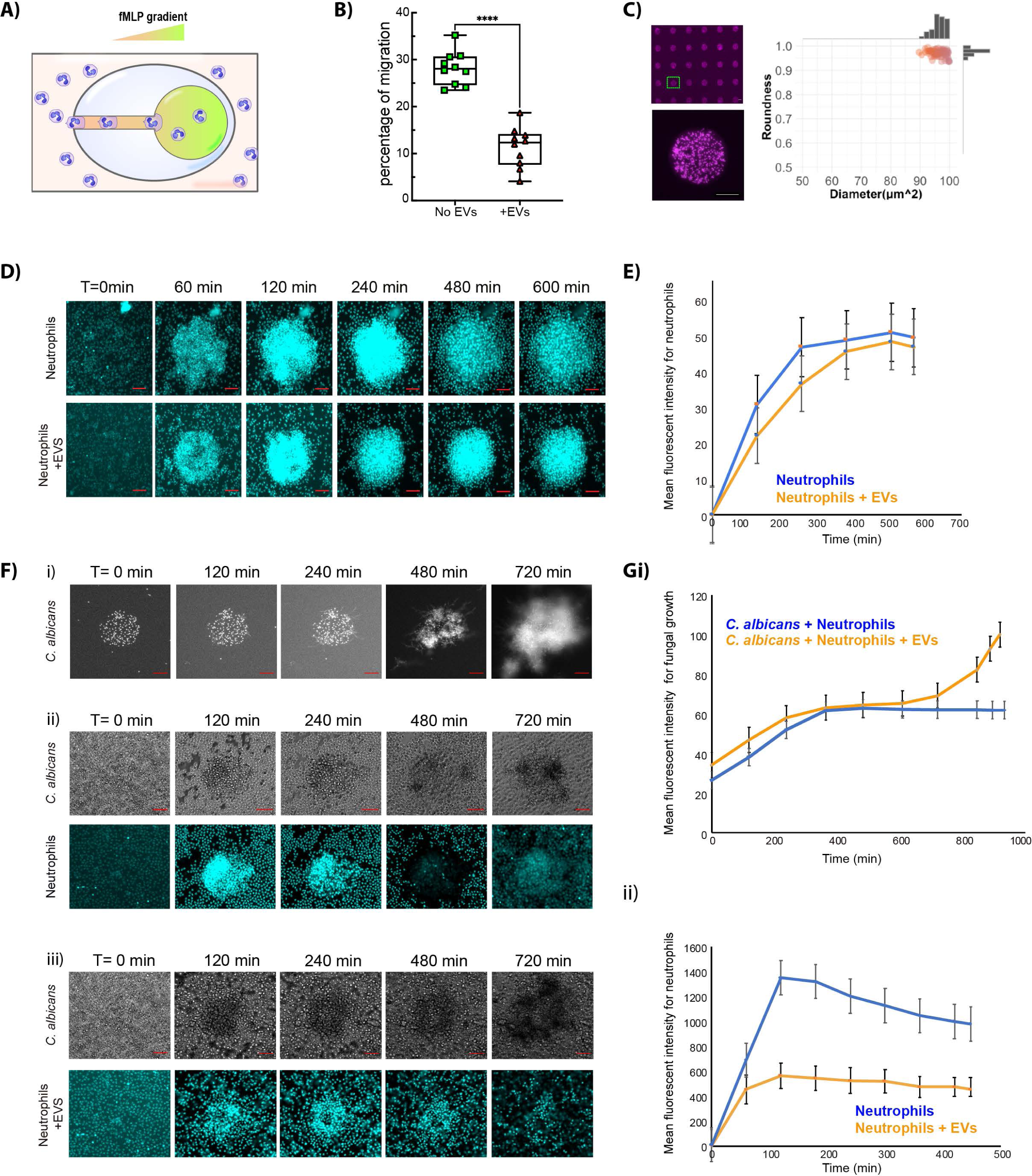
iRBC-EVs alters chemotaxis and swarming in primary neutrophils. **A)** Cartoon illustration of the chemotaxis egg shaped device (see method section for full description). **B)** The percentage of migration of untreated and iRBC-EVs treated primary neutrophils toward fMLP gradients using the egg-shaped device in A. **C)** Graph and images showing the uniformity in the size of each patterned *C. albicans* spot in the swarming assay device**. D)** Sequential images showing accumulation of untreated neutrophils and iRBC-EVs treated neutrophils at the swarm site at 0, 60, 120, 240, 480 and 600 minutes around zymosan particles (scale bar 25 μm). **E)** The dynamics of neutrophil swarm size in untreated neutrophils (blue line) and iRBC-EVs treated neutrophils (orange line). **Fi)** Sequential images showing *C. albicans* growth in absence of neutrophils at 0mins, 120mins, 240mins, 480mins and 720 mins (scale bar 100 μm). *C. albicans* expresses a far-red fluorescent protein. The intensity of the red dye is used to assess growth. Transformed black and white images are shown. **ii)** Sequential images showing the killing of C. *albicans* (far-red fluorescent protein) and swarm formation (blue, nuclei of neutrophils): untreated primary neutrophils. **iii)** iRBC-EVs treated primary neutrophils around *C. albicans* at 0mins, 120mins, 240mins, 480mins and 720 mins (Scale bar 100 μm, *n*=10 experiments). **Gi)** The growth dynamics of *C. albicans* with untreated primary neutrophils (blue line) and iRBC-EVs treated primary neutrophils (orange line). **ii)** The dynamics of primary neutrophils swarming. Untreated primary neutrophils (blue curve) and iRBC-EVs treated primary neutrophils (orange curve). N=60 swarms across three donors for figures E and G except for the control group. Error bars represent mean +/- standard error for these measurements.

For an effective microbe killing, neutrophils must exhibit a well-coordinated migration by swarming, thus we investigated the impact of iRBC-EV on swarming. The swarming assay device arranges microbes in clusters of 100-µm diameter, grouped in 8 × 8 arrays, separated inside individual wells, in a 16-well format (Figure 3C)(Babatunde et al., 2021). The device allows monitoring of the interaction of primary neutrophils and live *Candida albicans* or zymosan particles in real-time (Hopke et al., 2020).

First, we tested the effect of iRBC-EV on the swarming abilities of primary neutrophil and we observed that both iRBC-EV treated and untreated neutrophils swarm efficiently toward zymosan particle (dead yeast particles) (Figure 3D and 3E). The swarming curve shows that the mean fluorescent intensity for both groups increased sharply the first 60 minutes. However, the untreated neutrophil swarms increased significantly compared to iRBC-EV treated neutrophils, suggesting a larger swarm size in the untreated neutrophil (Figure 3D and 3E).

Next, to test the effect of iRBC-EV on the swarming abilities of primary neutrophils against *C. albicans* clusters (Figure 3F). We first monitor the growth of *C. albicans* growth in the absence of primary neutrophils in the swarming assay (Figure 3Fi, supplemental video 2). We observed that *C. albicans* clusters growth increased over time. In the presence of neutrophils, we observed that both iRBC-EV treated and untreated neutrophils swarm efficiently toward the *C. albicans* target [Figure 3F (ii & iii)]. Surprisingly, iRBC-EVs treated neutrophils also demonstrated a similar ability to arrest the growth of C. *albicans.* The growth curve shows that the mean fluorescence intensity for fungal growth for both groups increased sharply from ∼ 20 – 60 min. However, after ∼420 minutes, untreated but not iRBC-EV treated neutrophils were able to stop the growth of *C. albicans* [Figure 3G (i)] (Supplemental video 3). The mean fluorescent intensity for fungal growth of the iRBC-EVs treated neutrophils starts to peak after ∼420 minutes, from ∼60 - 100, suggesting reduced killing ability in neutrophil treated with iRBC-EVs [Figure 3G(i)] (Supplemental video 4). The mean swarm fluorescent intensity around *C. albicans* target was larger for untreated neutrophils compared to iRBC-EVs treated neutrophils, suggesting a larger swarm size around the microbial target in untreated compared to iRBC-EVs treated neutrophils [Figure 3G(ii)].

### Transcriptional Regulation of Inflammatory Response in primary neutrophils via EV

Neutrophil function is driven by complex signaling pathways that involve generating and receiving diverse signals from a multiplicity of sources, including other host cells and bacterial pathogens. To gain insight into the molecular processes that are modulated by EVs in human neutrophils, we used bulk RNA-seq to analyze global changes in the neutrophil transcriptome after iRBC-EV and iRBC-EV + lipopolysaccharide (LPS) stimulation (Figure 4A). We used four different conditions: untreated, EVs for 6 hours, LPS for 5 hours and finally the combination of EVs + LPS for 6 and 5 hours respectively. The PCA revealed donor-driven batch effect in the neutrophil (3 donors) that was corrected using ComBat-Seq on the count data. The ComBat-Seq corrected-PCA of gene expression patterns revealed four main clusters according to the different treatments (Figure 4B).

**Figure 4:**
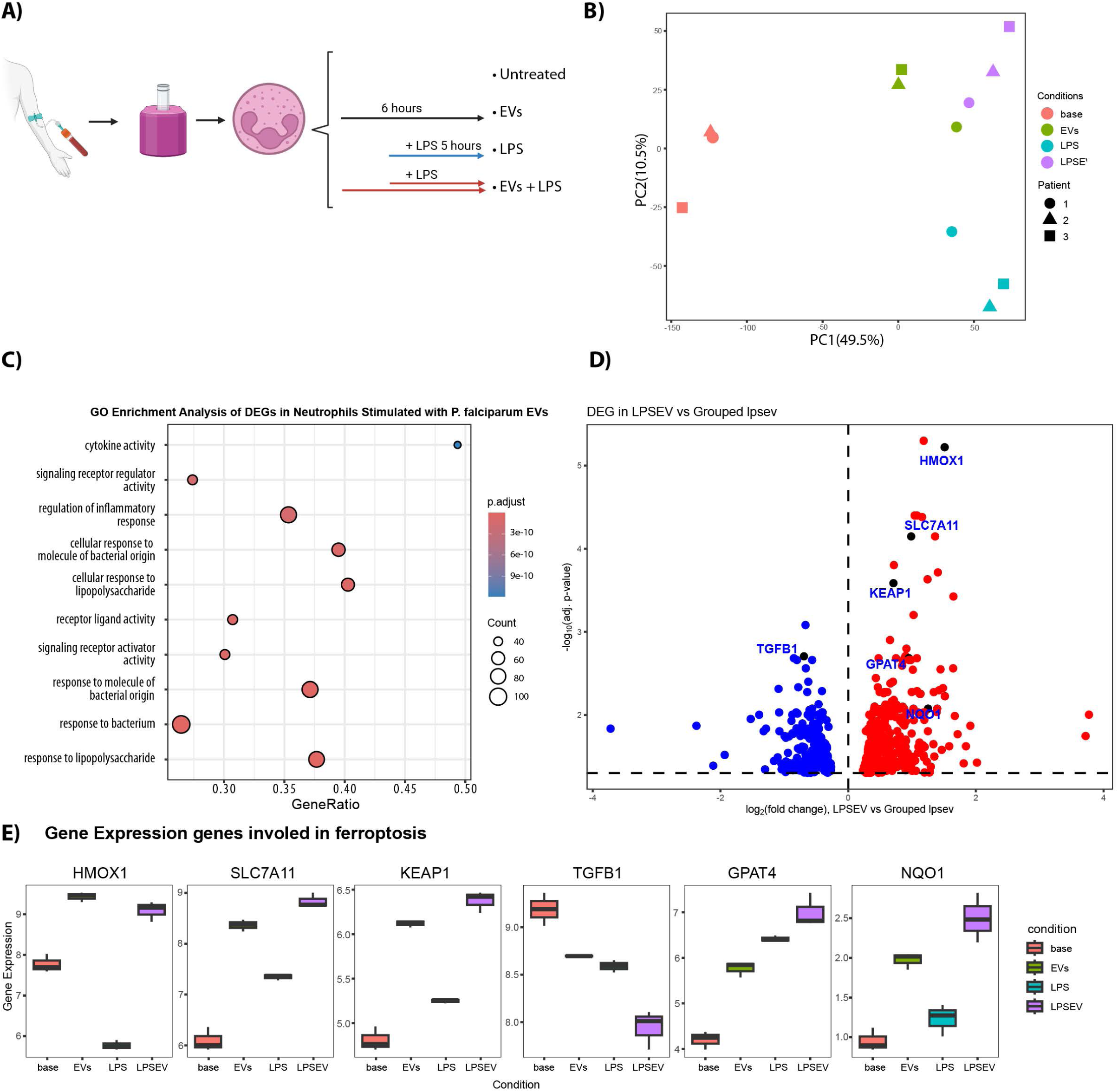
Transcriptional response of human neutrophils after LPS and EV stimulation. **A)** Experimental setup (n=3 for each stimulation). **B)** PCA of all samples using the full transcriptome. Shown in parentheses on the axes is the percentage of variance explained by each of the principal components. **C)** Enrichment analysis of DEGs in neutrophils stimulated with *P. falciparum* EVs compared to unstimulated neutrophils. Enrichment dot plot of the top Gene Ontology (GO) terms enriched among DEGs. Gene ratio: proportion of DEGs associated with each GO term relative to the total number of genes in that term. Enriched GO terms in y-axis are ordered by adjusted p-value (p.adjust). Dots size: number of DEGs contributing to each term. Dot color: adjusted p-value, with darker colors indicating more statistically significant terms. **D)** Volcano plot of DEGs in neutrophils stimulated with LPSEV compared to Grouped lpsev condition. All dots correspond to 71 genes with 20% change in expression ((LFC = log2(1.2). Significantly upregulated genes (red dots) and significantly downregulated genes (blues dots). The dashed horizontal line indicates the threshold for statistical significance (−log10(0.05)), and the vertical dashed lines represent the log2 fold change thresholds of 1.2 and −1.2. The labeled genes in blue and the corresponding black dot are genes involve in ferroptosis. **E)** Box plots display the expression levels of ferroptosis-related genes (HMOX1, KEAP1, NQO1, GPAT4, SLC7A11, and TGFB1) across different conditions (x-axis). The y-axis represents normalized gene expression values.

To find the genes differentially expressed in the LPS-EV but not in EV nor in LPS, we compared LPS-EVs condition against the combined group LPS + EV. This revealed 511 differentially expressed genes (DEGs), including 71 with 20% change in expression (Figure 4C) (Table S1)). Gene ontology enrichment showed that EV-treatment of neutrophil induces a molecular cascade of events that potentiate the expression of genes related to cytokine activity and regulation of both inflammatory responses and signaling receptor activity (Figure 4C). Interestingly, heme oxygenase 1 (HMOX1) was one the most up-regulated genes (Figure 4D) and given its role in ferroptosis, we analyzed if the DEGs in LPS-EV could be involved in ferroptosis and found five additional DEGs involved in ferroptosis (NQO1, SLC7AA11, KEAP1, GPAT4, TGFB1) (Figure 4D-E). Overall, our global analyses of the transcriptional landscape during iRBC-EV neutrophil interaction reveals a specific signature of the genes involved in ferroptosis in the presence of EVs.

### Transfer of miR451a to neutrophils by EVs modulates neutrophil function

Previously, we have demonstrated that miRNAs, and tRNAs-derived fragments were the most enriched small RNAs in EVs, with miR451a being the most abundant miRNA (Babatunde et al., 2018; Mantel et al., 2013). Given the efficient uptake of EVs by primary neutrophils, we hypothesized that EVs may deliver regulatory miRNAs to neutrophils. We incubated freshly isolated neutrophils and then determined miR451a levels in primary neutrophils via qPCR. We observed a 30-fold increase of miR451a expression in neutrophils upon EV treatment (Figure 5A). To determine whether miR451a originates from EV transfer, rather than being produced by neutrophils after EV uptake. We pre-treated neutrophils with α-amanitin before incubation with EVs (Lee et al., 2004). Since miR451a expression remains high even with treatment with α-amanitin, it shows that it is an actual transfer of miR451a to neutrophils by EVs (Figure 5A). Next, using RNA fluorescent *in situ* hybridization (FISH), we were able to directly detect and quantify miR451a copy numbers in primary neutrophils upon EV uptake. The miR451a uptake was significantly increased upon EV treatment (Figure 5B).

**Figure 5:**
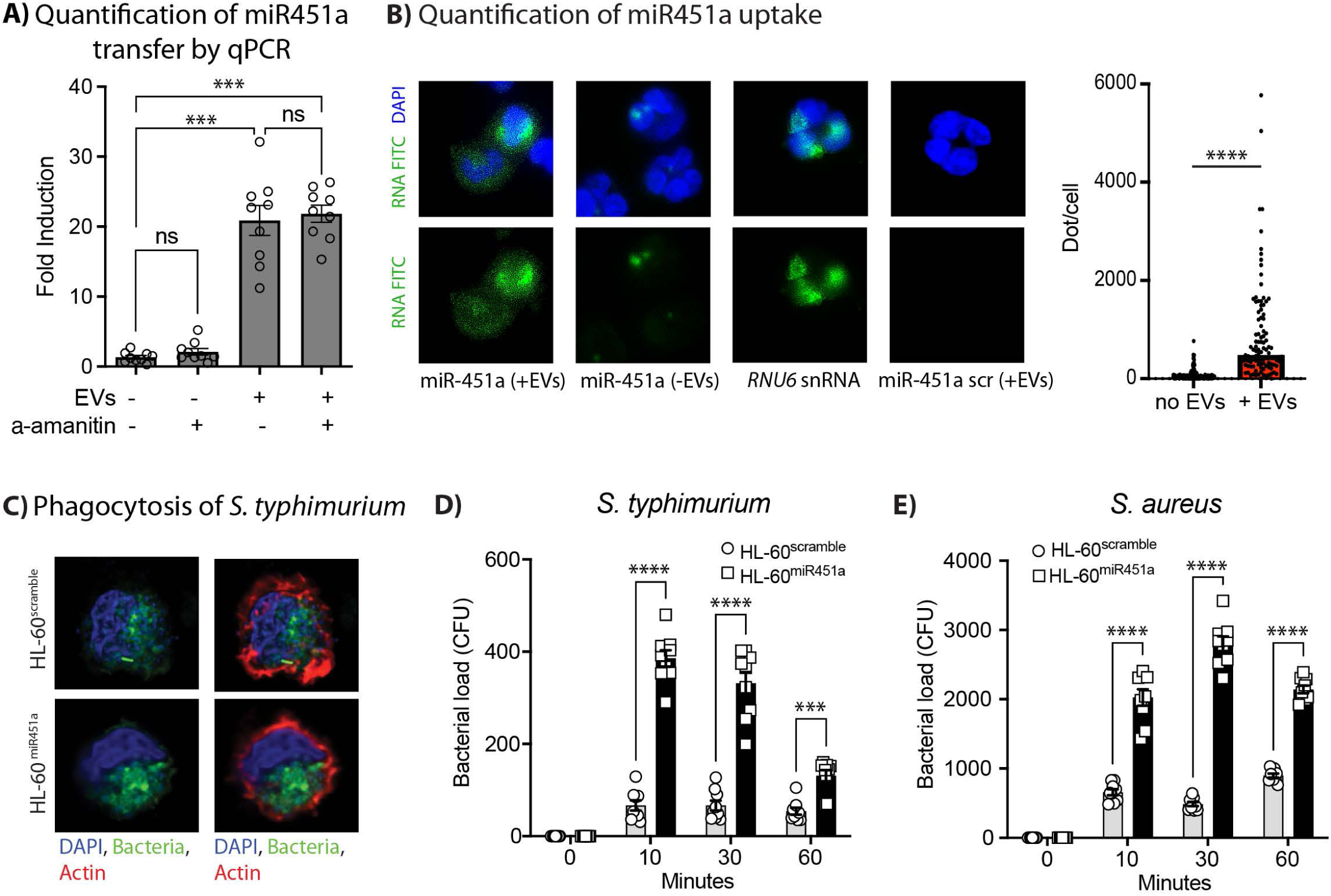
Transfer of miR451a to neutrophils by EVs and modulation of neutrophil function. A) EV uptake does not induce miRNA expression in neutrophils. Neutrophils, +/- pretreatment with α-amanitin for 30 minutes were incubated with EVs or left untreated. miR451a was quantified by qPCR in neutrophils upon EV incubation. qPCR results from all experiments are normalized by the 2^-Ct^ method, using *RNU6* as a reference and expressed as mean and fold induction over control (mean ± s.e.m.; *n*=3 experiments), *P* versus control (Student’s *t*-test). **B)** Detection of miRNA by RNA FISH. miRNA transfer to neutrophils by EVs is measured by RNA FISH. miR451a is only detected in neutrophils upon EV treatment, and only when a sequence-specific probe is used for detection (green spots: single-molecule RNA FISH; maximum intensity merges of Z-stacks). RNU6, positive control; scramble, negative control. Blue, nuclei (DAPI). P versus control (Student’s t-test). **C)** dHL60 expressing miR451 or scramble RNA were incubated with *Staphylococcus aureus* expressing GFP and phagocytosis was measured by fluorescent microscopy. **D)** To determine the antibacterial activity of dHL-60^miR451a^ and dHL-60^scramble^, the cells were incubated with *S. typhimurium* **E)** *S. aureus* in a CFU time course experiment. Viable bacteria colonies were counted after incubation (mean ± s.e.m.). N=9 from 3 independent experiments. Comparisons between dHL-60^miR451a^ and dHL-60^scramble^ at each time point were performed using a two-way ANOVA with the Sídák’s multiple comparisons test. Significance levels are indicated as \**P*<0.05, \*\**P*<0.01, \*\*\**P*<0.001 and \*\*\*\**P*<0.0001.

To ascertain the role of iRBC-EV derived miR451a in inhibiting ROS and antimicrobial functions in primary neutrophils, we transduced Human leukemia cell (HL-60 cells) with a lentiviral expression vector containing miR451a (or scramble control, shRNA against GFP). Next, we differentiated HL-60 cells into neutrophil-like cells via dimethyl sulfoxide (DMSO) treatment (Babatunde et al., 2021). We then compared the killing ability of HL-60 neutrophil-like cells expressing either miR451a (dHL-60^miR451a^) or a scramble dHL-60^scramble^ upon challenge with either *S. typhimurium* (Gram-negative) or *Staphylococcus aureus* (Gram-positive bacteria). Microscopy revealed that the capacity of HL-60^miR451a^ to subsequently ingest the bacteria remained unimpaired (Figure 5C). To test the killing ability of the HL-60 mutant cells, we challenged the cells with bacteria and monitored the killing for 60 minutes. The rate of killing in HL-60^miR451a^ was significantly impaired and slowed down compared to HL-60^scramble^ for both bacteria (Figure 5D-E). Importantly, we were able to recapitulate the observed effect of EVs on primary neutrophil functions in the HL-60^miR451a^ neutrophil-like cells, further suggesting a crucial role of iRBC-EV derived miR451a in the inhibition of the antimicrobial function of primary human neutrophils.

### miR451a alters chemotaxis and swarming in HL-60 neutrophil-like cells

To further investigate the role of EV derived miR451a in inhibiting chemotaxis and swarming, we tested the ability of HL-60^miR451a^ and HL-60^scramble^ to do chemotaxis and swarm using a tapered channel microfluidic device (Babatunde et al., 2021) and swarm assay respectively. The choice of a tapered channel device for this study was selected because it mimics the variation in blood vessel sizes and resistances encountered by neutrophils during migration. Using tapered-channel devices, we measured the fraction of HL-60^miR451a^ and HL-60^scramble^ cells migrating and quantified the frequency of various migration patterns. We found that 10% and 40% of HL-60^miR451a^ and HL-60^scramble^ cells migrated in N-Formyl-L-methionyl-L-leucyl-phenylalanine (fMLP) gradients, respectively (Figure 6A). Next, to quantify the frequency of previously reported neutrophil migration patterns in both mutant HL-60 cells. We focused on four migration patterns identified in earlier studies, including persistent migration (P), arrest (A), oscillation (O), and retro taxis (R)(Babatunde et al., 2021; Wang et al., 2018). Persistent migration indicates neutrophils that migrated through the channels without changing directions. Arrest describes neutrophils that are trapped in the channels. Oscillation indicates neutrophils that change migration direction more than two times. Retro-taxis describes neutrophils that migrated back to the cell-loading channel. We found that 70.9, 0.3, 20.9 and 7.9% of HL-60^scramble^ neutrophils, and 18.9, 1.9, 77.3 and 1.9% of HL-60^miR451a^ neutrophils like cells exhibited persistent, oscillation, arrest and retro taxis migratory patterns in fMLP gradients respectively (Figure 6B). To test the swarming ability of the HL-60 mutant strains, we use patterned zymosan clusters particles as microbial targets in the swarming assay. Three distinct phases of swarming have been previously reported (Lammermann et al., 2013). We observed that both HL-60^miR451a^ and HL-60^scramble^ neutrophils like cells swarm toward zymosan targets (Figure 6C). For the HL-60^scramble^ neutrophils like cells, swarming starts with random migration of the cells on the surface (*scouting phase*-5 min Figure 6C). After the first HL-60 neutrophil like cell interacts with the cluster, the number of migrating neutrophils towards the zymosan cluster increases rapidly (*growing phase-*Figure 6C). Swarms reach their peak size 220–240 min later, after which the size remains stable (*stabilization phase*-Figure 6C and 6D). In particular, the swarm area around the zymosan particle cluster for HL-60^scramble^ increased sharply from 0 to ∼ 35,000 μm^2^ in the first 240 min (*scouting and growing phase-*Fig. 6C and 6D) and after 240 minutes, the swarm size remained constant throughout the imaging time (*stabilization phase-* Figure 6C and 6D). We observed similar aggregation dynamics in HL-60^miR451a^ cells. Aggregation starts with the cells moving randomly on the surface of the zymosan particle clusters (*scouting phase*-5 min Figure 6C-*second panel*). Swarms in HL-60^miR451a^ cells reach their peak size after 100–120 min, after which the swarm size remained small and constant throughout the imaging time (*stabilization phase*-Figure 6C and 6D). The HL-60^miR451a^ swarm size increased from 0 to ∼ 12,000 μm^2^ in the first 2 hours (*growth phase*-Figure 6C and D) after which the swarm size remained small and constant (*stabilization phase*-Figure 6D). We found that the final mean swarm size of HL-60^scramble^ was 5 times larger as compared to HL-60^miR451a^, measuring 50,000 μm^2^ vs 10,000 μm^2^, respectively (Figure 6E).

**Figure 6:**
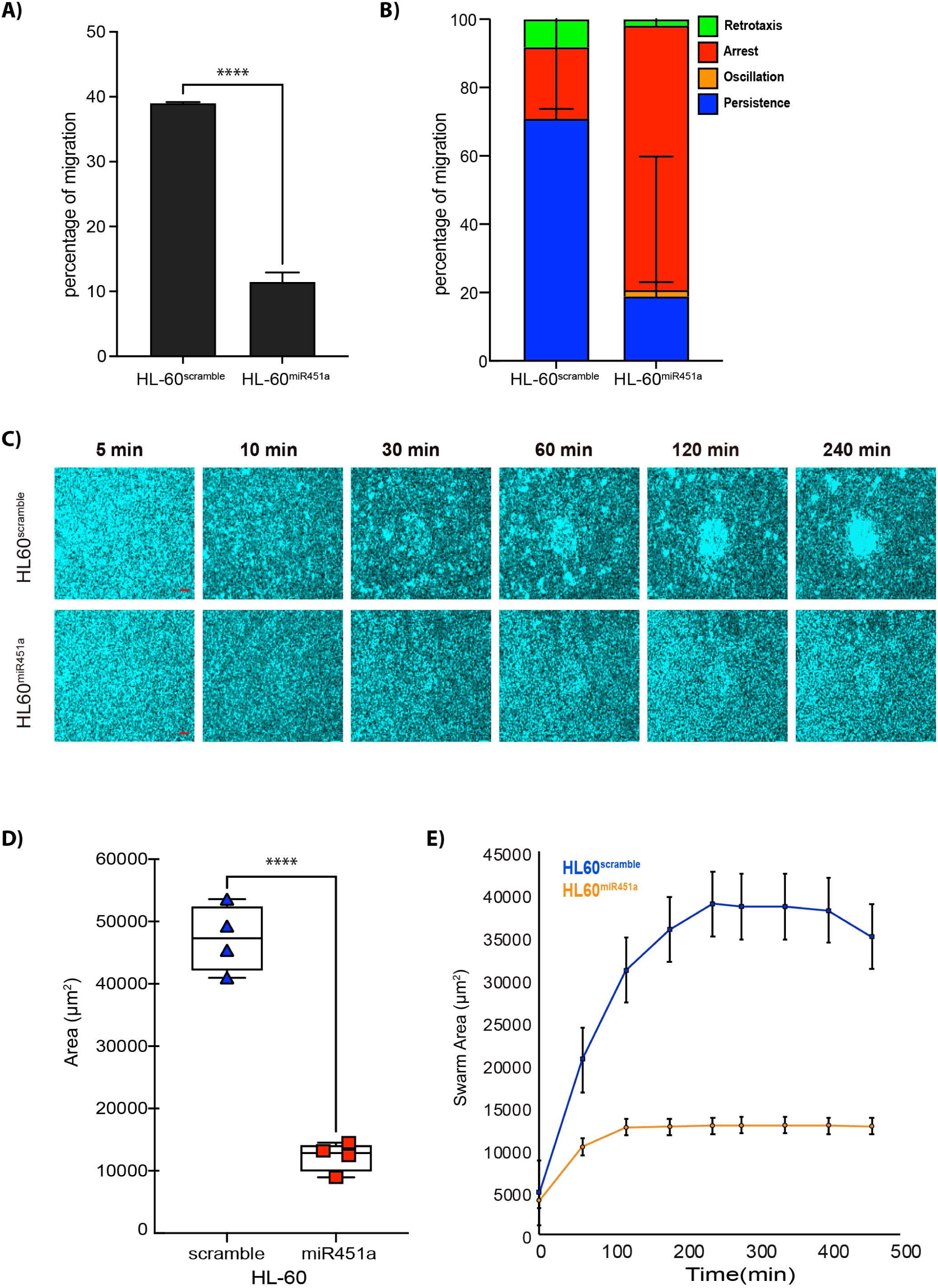
miR451a alters chemotaxis and swarming in differentiated HL-60 neutrophil like cells. **A)** The percentage of migration of differentiated HL-60^scramble^ and HL-60^miR451a^ toward fMLP gradients (mean ± s.e.m.; n=5 independent experiments). Comparisons were performed by unpaired, two-tailed t-test; \*\*\**P*<0.001. **B)** Percentage of migratory patterns in DdHL-60^scramble^ cells, ii) DdHL-60^miR451a^ cells. Persistence (Blue color), Arrest (Red color), Oscillation (Orange color) and Retrotaxis (Green color). **C)** Sequential images showing swarm formation in HL-60^scramble^ and HL-60^miR451a^ (blue) around patterned zymosan particle clusters at 5mins, 10mins, 30mins, 60mins, 120 mins and 240mins (scale bar: 25 µm). **D)** dHL-60^scramble^ (blue curve) accumulation on targets is fast and form a bigger swarm and dHL-60^miR451a^ (orange curve) aggregation proceeds fast and then continues slower over time. N=30 swarms and Error bars represent standard error for these measurements. Comparisons were performed by unpaired, two-tailed t-test; \*\*\*\**P*<0.0001. **E)** Comparison of swarm area formed between dHL-60^scramble^ and dHL-60^miR451a^. Comparisons were performed by unpaired, two-tailed t-test; \*\*\*\**P*<0.0001.

### miR451a modulates HL-60 neutrophil like cell transcriptional response to bacterial-like challenge

Next, to assess the effect of miR451a on gene expression, we performed RNA sequencing analysis on HL-60 human promyelocyte cells engineered to constitutively express miR451a via bulk and single cell RNA seq. For bulk RNA seq, we compared HL-60^scramble^ cells, HL-60^scramble^-LPS (90 mins), HL-60^scramble^ –LPS (180mins), HL-60^miR451a^, HL-60^miR451a^-LPS (90 mins) and HL-60^miR451a^-LPS (180 mins). We stimulated the cells for 90 and 180 mins to monitor gene regulation over time (Kobayashi et al., 2003). Differential gene expression analysis revealed a significantly higher number of DEGs (FDR < 0.05) in HL-60^miR451a^ cells compared to HL-60^scramble^ cells without LPS stimulation (46%). However, notably fewer significant DEGs were identified in the LPS-stimulated cells (Table S2). GO term enrichment analysis showed that miR451a induces a molecular cascade in HL-60 cells that involves 259 GO terms, and they are enriched with terms related to mitochondrial gene expression, translation and protein matrix in the absence of LPS stimulation (Figure 7A-B). Following 180 minutes of LPS stimulation, HL-60^miR451a^ cells exhibited significant enrichment in 126 GO terms compared to HL-60^scramble^ cells. Notably, the top 10 GO terms were linked to response to a molecule of bacterial origin such as LPS and cytokine activity and showed downregulation. These findings align with *in vitro* experiments, where genes associated with neutrophil functions like migration and chemotaxis, were downregulated in presence of EVs (Figure 7C-D). The PCA revealed distinct clustering patterns among the cell populations and treatment conditions. HL-60^scramble^ and HL-60^miR451a^ cells formed two separate clusters, indicating significant differences in their overall gene expression profiles. Furthermore, LPS activation for 90 and 180 minutes induced additional separation between these clusters. This time-dependent response to LPS stimulation was observed in both cell types, suggesting that the transcriptional changes elicited by LPS were substantial enough to be captured by PCA. The maintained separation between HL-60^scramble^ and HL-60^miR451a^ clusters following LPS treatment implies that the miR451a modification may alter the cellular response to inflammatory stimuli. These PCA results underscore the impact of miR451a on cellular phenotype and the dynamic nature of the cells’ response to LPS over time (Figure 7E). CNET analysis provided a comprehensive visualization of gene-pathway relationships, revealing that several enriched biological processes, including response to LPS, inflammatory response and positive regulation of ERK1 and ERK2 cascade, were driven by overlapping sets of genes (Figure 7F). The network structure identified TNF, RIPK2, IL-12B, as central nodes, linking multiple pathways. This suggests that miR451a genes may act as key regulators in coordinating the cellular response to bacterial infections.

**Figure 7:**
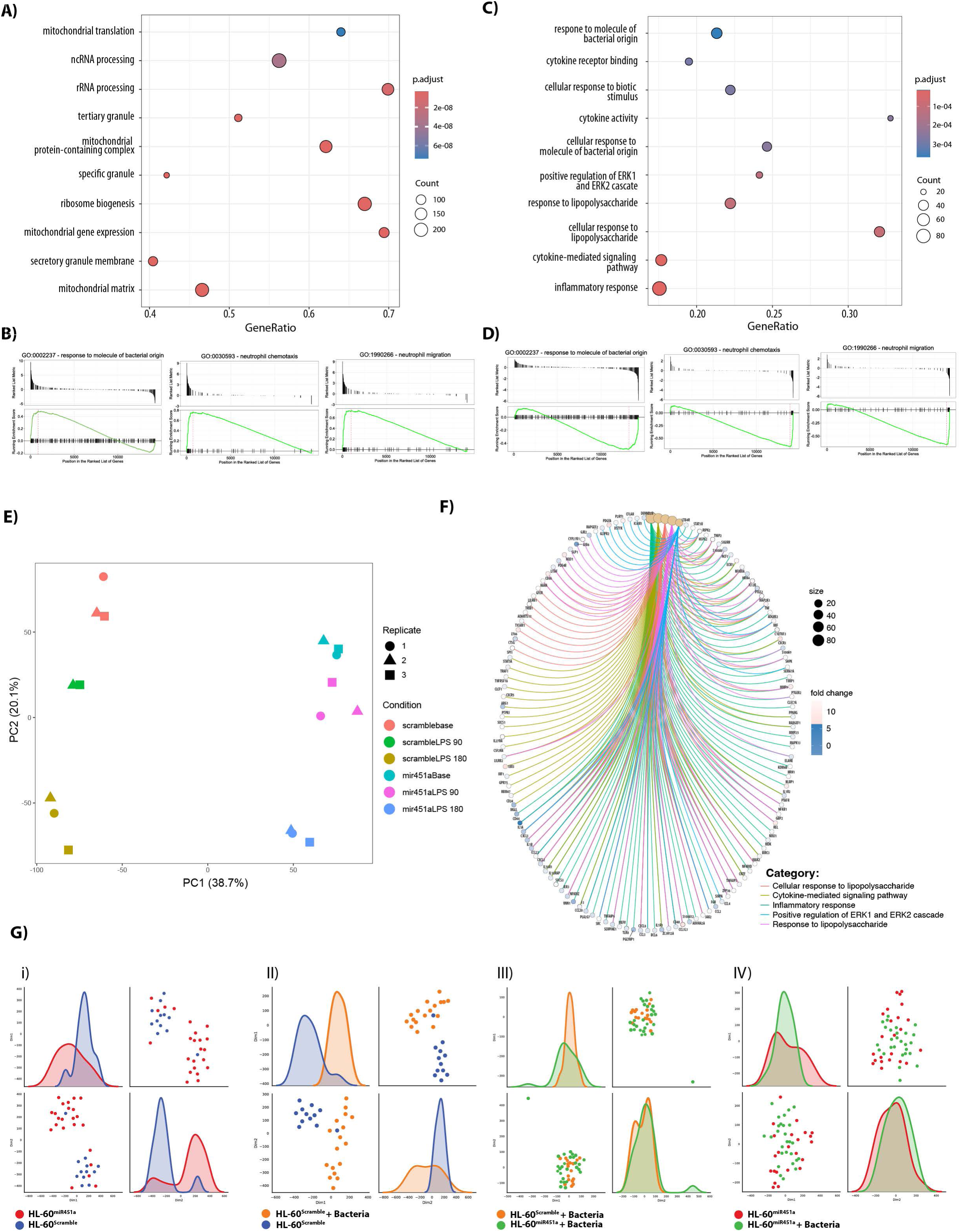
Enrichment analysis of DEGs in HL60^miR451a^ cells compared to HL60^scramble^ cells without LPS stimulation. **A, C)** Enrichment dot plot of the top Gene Ontology (GO) terms enriched among DEGs in HL-60^miR451a^ cells compared to HL-60^scramble^ cell without LPS (A) and in HL-60^miR451a^ cells compared to HL-60^scramble^ cells after LPS stimulation (C). Gene ratio indicates the proportion of differentially expressed genes associated with each GO term relative to the total genes annotated to that term. The enriched GO terms displayed on the y-axis are ranked according to their adjusted p-values (p.adjust). Dots size: number of DEGs contributing to each term. Dot color: adjusted p-value, with darker colors indicating more statistically significant terms. **B, D)** GSEA enrichment plot of the genes sets associated with three different GO terms in HL-60^miR451a^ cells compared to HL-60^scramble^ cell without LPS **B)** and in HL-60^miR451a^ cells compared to HL-60^scramble^ cells after LPS stimulation **D)**. **E)** Principal component analysis of all samples using the full transcriptome. **F)** Cnet plot of enriched biological pathways and associated genes. Circular nodes represent enriched pathways, while circles nodes represent genes. **G)** Figures highlighting pairwise PCA analysis of the main groups (HL-60^mir451a^ and HL-60^Scramble^, HL60^Scramble+Bacteria^ and HL-60^Scramble^, HL-60^Scramble+Bacteria^ and HL-60^miR451a+Bacteria^, HL-60^mir451a^ and HL-60^miR451a+Bacteria^. The projections are based on the top 1000 most variable genes and with the cells plotted on 1^st^ and 2^nd^ PCA components score. In each plot, a line distribution of the single cells within the PCA subspace is shown.

Next, we performed single cell RNA-seq to compare the control HL-60^scramble^ with the HL-60^miR451a^ cells and checked how they responded to bacterial exposure. We identified more than 1’000’000 read counts with 4’000 gene counts per cell types (Supplemental figure 1A-C). We identified several neutrophil genes expressed in the top 30 most expressed genes in all the cell populations (Supplemental figure 1D-H). The PCA of the differentially expressed genes (DEGs) clearly separated HL-60^scramble^ from the HL-60^miR451a^population (Figure 7Gi). when HL-60^scramble^ are activated with bacteria the population clearly separated (Figure 7Gii). In contrast, when HL-60^miR451a^ were activated with bacteria, they did not separate well from the unstimulated HL-60^miR451a^ (Figure 7Giii). To further characterize the transcriptional differences between cell types and conditions, we performed pairwise comparisons of gene expression profiles (Supplemental figure 2-5). Additionally, we examined the expression of published key markers associated with malaria and bacterial co-infection responses in HL-60^scramble^ and HL-60^miR451a^ cells exposed to bacteria. This analysis revealed a significantly higher expression of pro-inflammatory genes in HL-60^miR451a^-bacteria, including TNF, NCF1, NCF1B and NCF1C in HL-60^miR451a^-bacteria (Supplemental Figure 2-7B). Gene set enrichment analysis (GSEA) of DEGs between the HL-60^scramble^-bact and HL-60^miR451a^-bacteria revealed that genes related to biological processes and immunological signature (neutrophil migration, defense to Gram-bacteria, inflammatory response and cell killing) (Supplemental figure 6A-B).

## Discussion

Our findings reveal that miR451a-containing EVs from iRBCs contribute to neutrophil paralysis and dysfunction, highlighting a novel mechanism of immune evasion in malaria. The host’s survival during blood stage malaria hinges on two defense strategies: resistance, which aims to eliminate parasites, and tolerance, which prioritizes host fitness over pathogen clearance (McCarville and Ayres, 2018; Medzhitov et al., 2012). While tolerance may reduce self-inflicted damage, it potentially compromises resistance to secondary infections (Soares et al., 2017). Thus, neutrophil paralysis presents a double-edged sword: it benefits the parasite by reducing immune constraints yet potentially protecting the host from severe inflammatory damage. This delicate balance underscores the complex interplay between parasite virulence and host immune responses in determining malaria outcomes. Multiple factors released by iRBCs have been shown to promote tolerance by modulating the immune system, particularly through the suppression of neutrophil functions, such as heme (Cunnington et al., 2011) and food vacuoles (Dasari et al., 2012; Dasari et al., 2011). Here, we show that EV treatment impaired several aspects of neutrophil functionality, including ROS production, chemotaxis, anti-bacterial activity and swarming. The neutrophil “paralysis” appears to be part of the complex immune modulation induced by both *P. falciparum* and *P. vivax* infections. In malaria patients, neutrophils exhibit reduced chemotaxis and are a poor source of pro-inflammatory cytokines (Leoratti et al., 2012).

First, we confirmed that iRBC-EVs are efficiently internalized by neutrophils in an endocytic manner similar to endothelial cells (Mantel et al., 2013), in a process that requires actin polymerization (Dale et al., 2008). Coordinated neutrophil migration by swarming is crucial for arresting the growth and spread of various pathogens (Lammermann, 2016). We demonstrate that neutrophils-iRBC-EVs interaction alters the chemotactic response of human neutrophils towards fMLP, reduces their ability to swarm and to arrest the growth and progression of *C. albicans*. Additionally, these interactions trigger NETosis in both whole blood and isolated neutrophils *in vitro*. While EVs might contribute to NET release during malaria (Knackstedt et al., 2019), the ability of neutrophils to form large swarm is reduced, resulting in diminished antimicrobial activity at infection sites and increased pathogen proliferation.

Furthermore, we showed that EVs diminished the capacity of neutrophils to produce ROS upon PMA stimulation, consistent with observations of reduced ROS production in children with *P. falciparum* infection (Cunnington et al., 2012). Neutrophils treated with EVs also exhibit impaired killing of Gram-negative bacteria such as *Salmonella typhimurium*. these findings suggest that EVs contribute to the neutrophil dysfunction observed in malaria patients, potentially increasing the risk of death from bacteremia (Bachou et al., 2006; Bronzan et al., 2007; Mabey et al., 1987; Orf and Cunnington, 2015).

MiR451a is a microRNA critical for erythropoiesis (Chapman et al., 2017; Cheloufi et al., 2010; Murata et al., 2014; Patrick et al., 2010; Yu et al., 2010), is the most abundant miRNA in erythrocytes and regulates erythroblasts differentiation and oxidative stress (Kakumani et al., 2023). Given that EVs can shuttle functional RNA between cells (Mittelbrunn et al., 2011; O’Brien et al., 2020), we investigated whether miRNAs contained in EVs modulate neutrophil function. We focused on miR451a, due to its abundance in EVs (Babatunde et al., 2018). We demonstrated that EVs transferred miR451a to neutrophils. To further explore the molecular mechanisms, we generated a cell line overexpressing miR451a. HL-60^miR451a^ differentiated toward a neutrophil-like phenotype displayed the “paralyzed neutrophil” features, implicating miR451a directly in the suppression of chemotaxis, swarming and antibacterial activity. Previous studies have shown that miR451a has immunoregulatory properties, with reduced expression of miR451a observed in patients with rheumatoid arthritis and elevation of miR451a expression markedly reduces neutrophil chemotaxis (Murata et al., 2014). In addition to miRNAs, EVs from iRBCs can transfer parasitic genomic DNA to immune cells, activating STING-dependent DNA sensing and immune activation (Sisquella et al., 2017). While genomic DNA in EVs may activate monocytes, EVs also suppress CXCL10 secretion by disrupting ribosome association with CXCL10 transcripts, highlighting the complex nature of EV-mediated immune modulation (Ofir-Birin et al., 2021).

We next examined the effect of EVs on neutrophil transcriptional activity, particularly in response to bacterial stimulation. The transcriptomics data revealed that HO-1 is the most upregulated gene following EV treatment, whereas it is slightly downregulated by LPS alone. HO-1, encoded by HMOX1, plays a key role in malaria (Cunnington et al., 2012; Silva et al., 2020), with promoter polymorphisms and high HO-1 levels associated with severe malaria (Walther et al., 2012). Evidence from the rodent model showed that HO-1 plays a protective role by inducing tolerance and reducing damage to the host, since mice deficient in HO-1 are more susceptible to cerebral malaria (Ferreira et al., 2011; Pamplona et al., 2007).

HO-1 is crucial role for heme catabolism and provides anti-inflammatory, antioxidant, cytoprotective and vascular protection. Moreover, HMOX1 plays a complex and significant role in ferroptosis a form of programmed cell death characterized by iron-dependent lipid peroxidation (Jiang et al., 2021). We found that several genes involved in ferroptosis were upregulated, neutrophils appear particularly sensitive to ferroptosis, which can promote tolerance and tumor growth by limiting antitumor activity (Kim et al., 2022).

Neutrophils are crucial players in the immune response to malaria, but their activation must be carefully regulated. While they contribute to parasite clearance, excessive neutrophil activation and NET formation can lead to tissue damage and exacerbate disease severity. Future research and therapeutic strategies may focus on harnessing the protective functions of neutrophils while mitigating their potential harmful effects in severe malaria.

Collectively, our *in vitro* data suggest that iRBC-EVs released during malaria contribute to the host immune suppression and increased susceptibility to opportunistic infections by inhibiting neutrophil functions such as swarming and antimicrobial activity. Previous observations of elevated iRBC-EVs in severe malaria patients (Nantakomol et al., 2011) strongly support a role for these vesicles in malaria pathology.

In summary, our study identifies miR451a-containing EVs from iRBCs as key mediators of neutrophil dysfunction, providing new insights into how malaria parasites subvert host immunity. By impairing critical neutrophil functions such as chemotaxis, swarming, and antimicrobial activity, these EVs contribute to both immune evasion and increased susceptibility to secondary infections. This dual effect underscores the complex evolutionary balance between parasite survival strategies and host defense mechanisms, where immune modulation may simultaneously limit immunopathology and compromise resistance (Su et al., 2025). Our findings highlight the importance of targeting EV-mediated pathways in future therapeutic strategies aimed at restoring neutrophil function and improving outcomes in malaria.

## Acknowledgements

This work was partially supported by the Swiss National Science Foundation (grant# 31003A_182729 and grant# CRSK-3_190848 to PYM), the Novartis foundation for medical-biological research (P.-Y.M.), he Kurt and Senta Herrmann Foundation, the Gottfried and Julia Bangerter-Rhyner-Stiftung (PYM) and the Research Pool of the University of Fribourg. Additional grants include the Swiss Excellence Scholarships for Foreign Scholars (grant # 2016.0934 to K.A.B), Swiss National Science Foundation mobility (grant # P1FRP3_181378 and P500PB_203002 to K.A.B) and Jubiläumsstiftung von Swiss Life (grant # 01-1242, 01-1309 and 01-1350 to K.A.B)

## Authorship contributions

P.-Y.M., conceived and conceptualized the study. P-Y.M., F.S., R.S., M.H.-J. L.F., M.W., D.I., provided the methodology by establishing all major assays. K.A.B. M.N., P.M.M, B.S., A.H., S.L., M.N., G.M., M.H-J., conducted the investigation, performed the experiments, and analyzed the data. P.-Y.M. and K.A.B. wrote the original draft of the manuscript. P.M.M and M.W. wrote, reviewed, and edited the manuscript.

## Disclosure of Conflicts of Interest

The authors declare no competing interests.

## Declaration of generative AI and AI-assisted technologies

During the writing of this manuscripts the authors used ChatGPT and Perplexity in order to improve the readability and language of the manuscript. After using this tool, the authors reviewed and edited the content as needed and take full responsibility for the content of the publication.

## Declaration of interests

The authors declare that they have no conflicts of interest.

## Resource availability

Lead contact

Further information and requests for resources and reagents should be directed to and will be fulfilled by the lead contact, Pierre-Yves Mantel.

## Supplemental information

Document S1. Figures S1–S6, Table S1 and Table S2.

Video S1.

Video S2

Video S3

Video S4

## Notes

### Competing Interest Statement

The authors have declared no competing interest.

## REFERENCES

Babatunde, K.A., Mbagwu, S., Hernandez-Castaneda, M.A., Adapa, S.R., Walch, M., Filgueira, L., Falquet, L., Jiang, R.H.Y., Ghiran, I., and Mantel, P.Y. (2018). Malaria infected red blood cells release small regulatory RNAs through extracellular vesicles. Sci Rep 8, 884.

Babatunde, K.A., Wang, X., Hopke, A., Lannes, N., Mantel, P.Y., and Irimia, D. (2021). Chemotaxis and swarming in differentiated HL-60 neutrophil-like cells. Sci Rep 11, 778.

Babatunde, K.A., Yesodha Subramanian, B., Ahouidi, A.D., Martinez Murillo, P., Walch, M., and Mantel, P.Y. (2020). Role of Extracellular Vesicles in Cellular Cross Talk in Malaria. Front Immunol 11, 22.

Bachou, H., Tylleskar, T., Kaddu-Mulindwa, D.H., and Tumwine, J.K. (2006). Bacteraemia among severely malnourished children infected and uninfected with the human immunodeficiency virus-1 in Kampala, Uganda. BMC infectious diseases 6, 160.

Brinkmann, V., Laube, B., Abu Abed, U., Goosmann, C., and Zychlinsky, A. (2010). Neutrophil extracellular traps: how to generate and visualize them. J Vis Exp.

Bronzan, R.N., Taylor, T.E., Mwenechanya, J., Tembo, M., Kayira, K., Bwanaisa, L., Njobvu, A., Kondowe, W., Chalira, C., Walsh, A.L., et al. (2007). Bacteremia in Malawian children with severe malaria: prevalence, etiology, HIV coinfection, and outcome. J Infect Dis 195, 895–904.

Chapman, L.M., Ture, S.K., Field, D.J., and Morrell, C.N. (2017). miR-451 limits CD4(+) T cell proliferative responses to infection in mice. Immunol Res 65, 828–840.

Cheloufi, S., Dos Santos, C.O., Chong, M.M., and Hannon, G.J. (2010). A dicer-independent miRNA biogenesis pathway that requires Ago catalysis. Nature 465, 584–589.

Cunnington, A.J., de Souza, J.B., Walther, M., and Riley, E.M. (2011). Malaria impairs resistance to Salmonella through heme- and heme oxygenase-dependent dysfunctional granulocyte mobilization. Nat Med 18, 120–127.

Cunnington, A.J., Njie, M., Correa, S., Takem, E.N., Riley, E.M., and Walther, M. (2012). Prolonged neutrophil dysfunction after Plasmodium falciparum malaria is related to hemolysis and heme oxygenase-1 induction. J Immunol 189, 5336–5346.

Dabritz, J., Schneider, M., Just-Nuebling, G., and Groll, A.H. (2011). Minireview: Invasive fungal infection complicating acute Plasmodium falciparum malaria. Mycoses 54, 311–317.

Dale, D.C., Boxer, L., and Liles, W.C. (2008). The phagocytes: neutrophils and monocytes. Blood 112, 935–945.

Dasari, P., Heber, S.D., Beisele, M., Torzewski, M., Reifenberg, K., Orning, C., Fries, A., Zapf, A.L., Baumeister, S., Lingelbach, K., et al. (2012). Digestive vacuole of Plasmodium falciparum released during erythrocyte rupture dually activates complement and coagulation. Blood 119, 4301–4310.

Dasari, P., Reiss, K., Lingelbach, K., Baumeister, S., Lucius, R., Udomsangpetch, R., Bhakdi, S.C., and Bhakdi, S. (2011). Digestive vacuoles of Plasmodium falciparum are selectively phagocytosed by and impair killing function of polymorphonuclear leukocytes. Blood 118, 4946–4956.

Dobin, A., Davis, C.A., Schlesinger, F., Drenkow, J., Zaleski, C., Jha, S., Batut, P., Chaisson, M., and Gingeras, T.R. (2013). STAR: ultrafast universal RNA-seq aligner. Bioinformatics 29, 15–21.

Eckerle, I., Ebinger, D., Gotthardt, D., Eberhardt, R., Schnabel, P.A., Stremmel, W., Junghanss, T., and Eisenbach, C. (2009). Invasive Aspergillus fumigatus infection after Plasmodium falciparum malaria in an immuno-competent host: case report and review of literature. Malar J 8, 167.

Ellett, F., Jalali, F., Marand, A.L., Jorgensen, J., Mutlu, B.R., Lee, J., Raff, A.B., and Irimia, D. (2019). Microfluidic arenas for war games between neutrophils and microbes. Lab Chip 19, 1205–1216.

Ferreira, A., Marguti, I., Bechmann, I., Jeney, V., Chora, A., Palha, N.R., Rebelo, S., Henri, A., Beuzard, Y., and Soares, M.P. (2011). Sickle hemoglobin confers tolerance to Plasmodium infection. Cell 145, 398–409.

Hocqueloux, L., Bruneel, F., Pages, C.L., and Vachon, F. (2000). Fatal invasive aspergillosis complicating severe Plasmodium falciparum malaria. Clinical infectious diseases: an official publication of the Infectious Diseases Society of America 30, 940–942.

Hopke, A., Nicke, N., Hidu, E.E., Degani, G., Popolo, L., and Wheeler, R.T. (2016). Neutrophil Attack Triggers Extracellular Trap-Dependent Candida Cell Wall Remodeling and Altered Immune Recognition. PLoS Pathog 12, e1005644.

Hopke, A., Scherer, A., Kreuzburg, S., Abers, M.S., Zerbe, C.S., Dinauer, M.C., Mansour, M.K., and Irimia, D. (2020). Neutrophil swarming delays the growth of clusters of pathogenic fungi. Nat Commun 11, 2031.

Jiang, X., Stockwell, B.R., and Conrad, M. (2021). Ferroptosis: mechanisms, biology and role in disease. Nat Rev Mol Cell Biol 22, 266–282.

Kakumani, P.K., Ko, Y., Ramakrishna, S., Christopher, G., Dodgson, M., Shrinet, J., Harvey, L.M., Shin, C., and Simard, M.J. (2023). CSDE1 promotes miR-451 biogenesis. Nucleic Acids Res 51, 9385–9396.

Kim, R., Hashimoto, A., Markosyan, N., Tyurin, V.A., Tyurina, Y.Y., Kar, G., Fu, S., Sehgal, M., Garcia-Gerique, L., Kossenkov, A., et al. (2022). Ferroptosis of tumour neutrophils causes immune suppression in cancer. Nature 612, 338–346.

Knackstedt, S.L., Georgiadou, A., Apel, F., Abu-Abed, U., Moxon, C.A., Cunnington, A.J., Raupach, B., Cunningham, D., Langhorne, J., Kruger, R., et al. (2019). Neutrophil extracellular traps drive inflammatory pathogenesis in malaria. Sci Immunol 4.

Kobayashi, S.D., Braughton, K.R., Whitney, A.R., Voyich, J.M., Schwan, T.G., Musser, J.M., and DeLeo, F.R. (2003). Bacterial pathogens modulate an apoptosis differentiation program in human neutrophils. Proc Natl Acad Sci U S A 100, 10948–10953.

Kotepui, M., Phunphuech, B., Phiwklam, N., Chupeerach, C., and Duangmano, S. (2014). Effect of malarial infection on haematological parameters in population near Thailand-Myanmar border. Malar J 13, 218.

Lamaze, C., Fujimoto, L.M., Yin, H.L., and Schmid, S.L. (1997). The actin cytoskeleton is required for receptor-mediated endocytosis in mammalian cells. J Biol Chem 272, 20332–20335.

Lammermann, T. (2016). In the eye of the neutrophil swarm-navigation signals that bring neutrophils together in inflamed and infected tissues. J Leukoc Biol 100, 55–63.

Lammermann, T., Afonso, P.V., Angermann, B.R., Wang, J.M., Kastenmuller, W., Parent, C.A., and Germain, R.N. (2013). Neutrophil swarms require LTB4 and integrins at sites of cell death in vivo. Nature 498, 371–375.

Lee, Y., Kim, M., Han, J., Yeom, K.H., Lee, S., Baek, S.H., and Kim, V.N. (2004). MicroRNA genes are transcribed by RNA polymerase II. EMBO J 23, 4051–4060.

Leoratti, F.M., Trevelin, S.C., Cunha, F.Q., Rocha, B.C., Costa, P.A., Gravina, H.D., Tada, M.S., Pereira, D.B., Golenbock, D.T., Antonelli, L.R., et al. (2012). Neutrophil paralysis in Plasmodium vivax malaria. PLoS Negl Trop Dis 6, e1710.

Lu, J., and Tsourkas, A. (2011). Quantification of miRNA abundance in single cells using locked nucleic acid-FISH and enzyme-labeled fluorescence. Methods Mol Biol 680, 77–88.

Mabey, D.C., Brown, A., and Greenwood, B.M. (1987). Plasmodium falciparum malaria and Salmonella infections in Gambian children. J Infect Dis 155, 1319–1321.

Mantel, P.Y., Hoang, A.N., Goldowitz, I., Potashnikova, D., Hamza, B., Vorobjev, I., Ghiran, I., Toner, M., Irimia, D., Ivanov, A.R., et al. (2013). Malaria-Infected Erythrocyte-Derived Microvesicles Mediate Cellular Communication within the Parasite Population and with the Host Immune System. Cell Host Microbe 13, 521–534.

McCarville, J.L., and Ayres, J.S. (2018). Disease tolerance: concept and mechanisms. Curr Opin Immunol 50, 88–93.

Medzhitov, R., Schneider, D.S., and Soares, M.P. (2012). Disease tolerance as a defense strategy. Science 335, 936–941.

Mittelbrunn, M., Gutiérrez-Vázquez, C., Villarroya-Beltri, C., González, S., Sánchez-Cabo, F., González, M.Á., Bernad, A., and Sánchez-Madrid, F. (2011). Unidirectional transfer of microRNA-loaded exosomes from T cells to antigen-presenting cells. Nature communications 2, 282.

Murata, K., Yoshitomi, H., Furu, M., Ishikawa, M., Shibuya, H., Ito, H., and Matsuda, S. (2014). MicroRNA-451 down-regulates neutrophil chemotaxis via p38 MAPK. Arthritis Rheumatol 66, 549–559.

Nantakomol, D., Dondorp, A.M., Krudsood, S., Udomsangpetch, R., Pattanapanyasat, K., Combes, V., Grau, G.E., White, N.J., Viriyavejakul, P., Day, N.P., et al. (2011). Circulating red cell-derived microparticles in human malaria. J Infect Dis 203, 700–706.

O’Brien, K., Breyne, K., Ughetto, S., Laurent, L.C., and Breakefield, X.O. (2020). RNA delivery by extracellular vesicles in mammalian cells and its applications. Nat Rev Mol Cell Biol 21, 585–606.

Ofir-Birin, Y., Ben Ami Pilo, H., Cruz Camacho, A., Rudik, A., Rivkin, A., Revach, O.Y., Nir, N., Block Tamin, T., Abou Karam, P., Kiper, E., et al. (2021). Malaria parasites both repress host CXCL10 and use it as a cue for growth acceleration. Nat Commun 12, 4851.

Orf, K., and Cunnington, A.J. (2015). Infection-related hemolysis and susceptibility to Gram-negative bacterial co-infection. Front Microbiol 6, 666.

Organization., W.H. (2022). World malaria report. Geneva World Health Organization.

Pamplona, A., Ferreira, A., Balla, J., Jeney, V., Balla, G., Epiphanio, S., Chora, A., Rodrigues, C.D., Gregoire, I.P., Cunha-Rodrigues, M., et al. (2007). Heme oxygenase-1 and carbon monoxide suppress the pathogenesis of experimental cerebral malaria. Nat Med 13, 703–710.

Patrick, D.M., Zhang, C.C., Tao, Y., Yao, H., Qi, X., Schwartz, R.J., Jun-Shen Huang, L., and Olson, E.N. (2010). Defective erythroid differentiation in miR-451 mutant mice mediated by 14-3-3zeta. Genes & development 24, 1614–1619.

Picelli, S., Bjorklund, A.K., Faridani, O.R., Sagasser, S., Winberg, G., and Sandberg, R. (2013). Smart-seq2 for sensitive full-length transcriptome profiling in single cells. Nat Methods 10, 1096–1098.

Regev-Rudzki, N., Wilson, D.W., Carvalho, T.G., Sisquella, X., Coleman, B.M., Rug, M., Bursac, D., Angrisano, F., Gee, M., Hill, A.F., et al. (2013). Cell-cell communication between malaria-infected red blood cells via exosome-like vesicles. Cell 153, 1120–1133.

Robinson, M.D., McCarthy, D.J., and Smyth, G.K. (2010). edgeR: a Bioconductor package for differential expression analysis of digital gene expression data. Bioinformatics 26, 139–140.

Ruhnke, M., Eichenauer, E., Searle, J., and Lippek, F. (2000). Fulminant tracheobronchial and pulmonary aspergillosis complicating imported Plasmodium falciparum malaria in an apparently immunocompetent woman. Clinical infectious diseases: an official publication of the Infectious Diseases Society of America 30, 938–940.

Silva, R., Travassos, L.H., Paiva, C.N., and Bozza, M.T. (2020). Heme oxygenase-1 in protozoan infections: A tale of resistance and disease tolerance. PLoS Pathog 16, e1008599.

Sisquella, X., Ofir-Birin, Y., Pimentel, M.A., Cheng, L., Abou Karam, P., Sampaio, N.G., Penington, J.S., Connolly, D., Giladi, T., Scicluna, B.J., et al. (2017). Malaria parasite DNA-harbouring vesicles activate cytosolic immune sensors. Nat Commun 8, 1985.

Snow, R.W., Guerra, C.A., Noor, A.M., Myint, H.Y., and Hay, S.I. (2005). The global distribution of clinical episodes of Plasmodium falciparum malaria. Nature 434, 214–217.

Soares, M.P., Teixeira, L., and Moita, L.F. (2017). Disease tolerance and immunity in host protection against infection. Nat Rev Immunol 17, 83–96.

Su, X.Z., Xu, F., Stadler, R.V., Teklemichael, A.A., and Wu, J. (2025). Malaria: Factors affecting disease severity, immune evasion mechanisms, and reversal of immune inhibition to enhance vaccine efficacy. PLoS Pathog 21, e1012853.

Vasquez, R.J., Howell, B., Yvon, A.M., Wadsworth, P., and Cassimeris, L. (1997). Nanomolar concentrations of nocodazole alter microtubule dynamic instability in vivo and in vitro. Mol Biol Cell 8, 973–985.

Venkatesan, P. (2024). The 2023 WHO World malaria report. Lancet Microbe 5, e214.

Walther, M., De Caul, A., Aka, P., Njie, M., Amambua-Ngwa, A., Walther, B., Predazzi, I.M., Cunnington, A., Deininger, S., Takem, E.N., et al. (2012). HMOX1 gene promoter alleles and high HO-1 levels are associated with severe malaria in Gambian children. PLoS Pathog 8, e1002579.

Wang, X., Jodoin, E., Jorgensen, J., Lee, J., Markmann, J.J., Cataltepe, S., and Irimia, D. (2018). Progressive mechanical confinement of chemotactic neutrophils induces arrest, oscillations, and retrotaxis. J Leukoc Biol 104, 1253–1261.

Were, T., Davenport, G.C., Hittner, J.B., Ouma, C., Vulule, J.M., Ong’echa, J.M., and Perkins, D.J. (2011). Bacteremia in Kenyan children presenting with malaria. J Clin Microbiol 49, 671–676.

Wilson, A.P., Wright, S., and Bellingan, G. (2000). Disseminated fungal infection following falciparum malaria. J Infect 40, 202–204.

Wolf, F.A., Angerer, P., and Theis, F.J. (2018). SCANPY: large-scale single-cell gene expression data analysis. Genome Biol 19, 15.

Yu, D., dos Santos, C.O., Zhao, G., Jiang, J., Amigo, J.D., Khandros, E., Dore, L.C., Yao, Y., D’Souza, J., Zhang, Z., et al. (2010). miR-451 protects against erythroid oxidant stress by repressing 14-3-3zeta. Genes & development 24, 1620–1633.

Yu, G., Wang, L.G., Han, Y., and He, Q.Y. (2012). clusterProfiler: an R package for comparing biological themes among gene clusters. OMICS 16, 284–287.

