## Supplementary Data for "Plasmodium falciparum subverts neutrophil function via host miR-451a loaded extracellular vesicles driving bacterial superinfection susceptibility"

### Supplementary Figure 1. Dataset overview

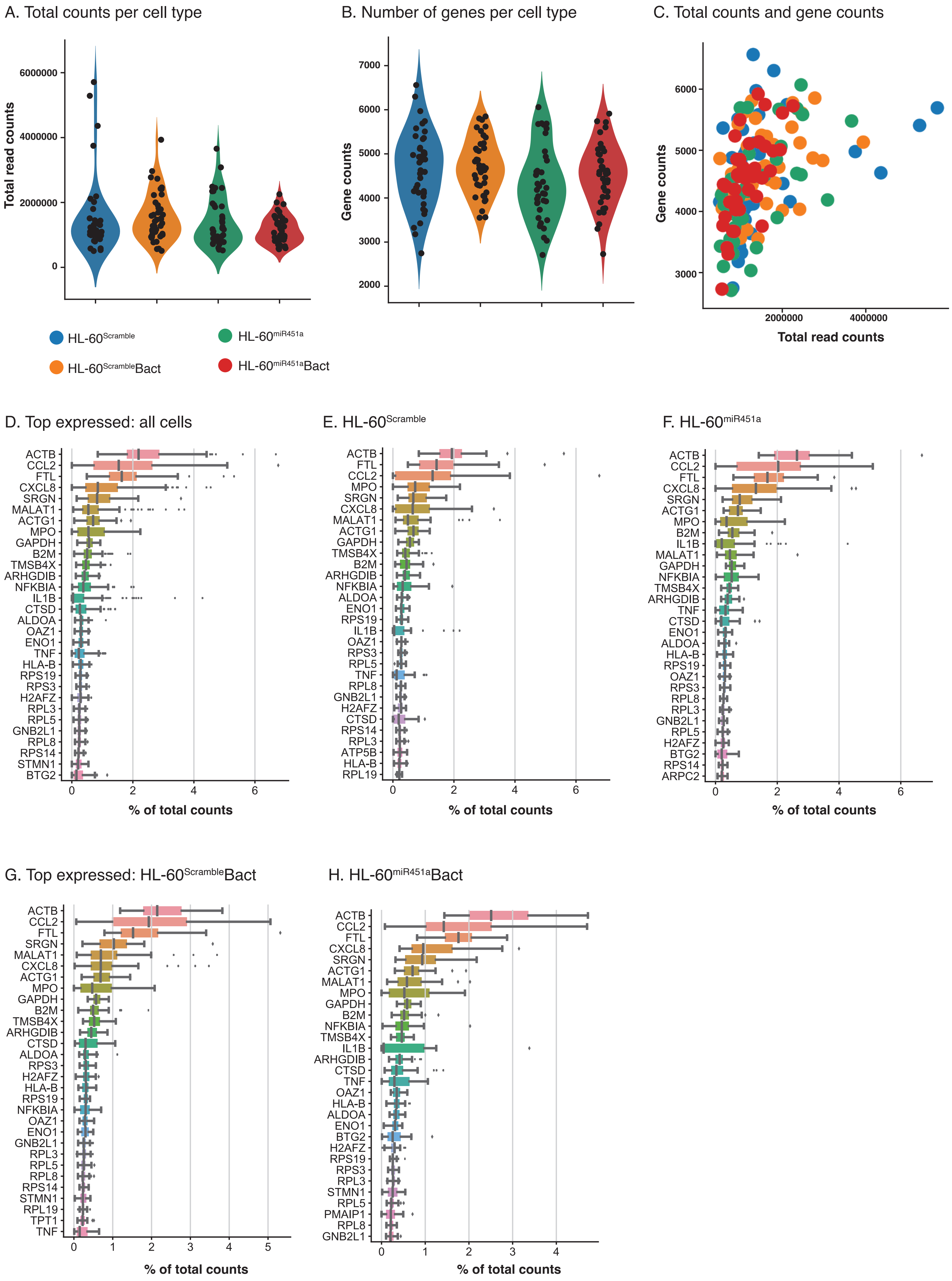

**Supplementary Figure 2. Cell types pairwise analysis (HL-60$^{scramble}$ and HL-60$^{mir451a}$)**

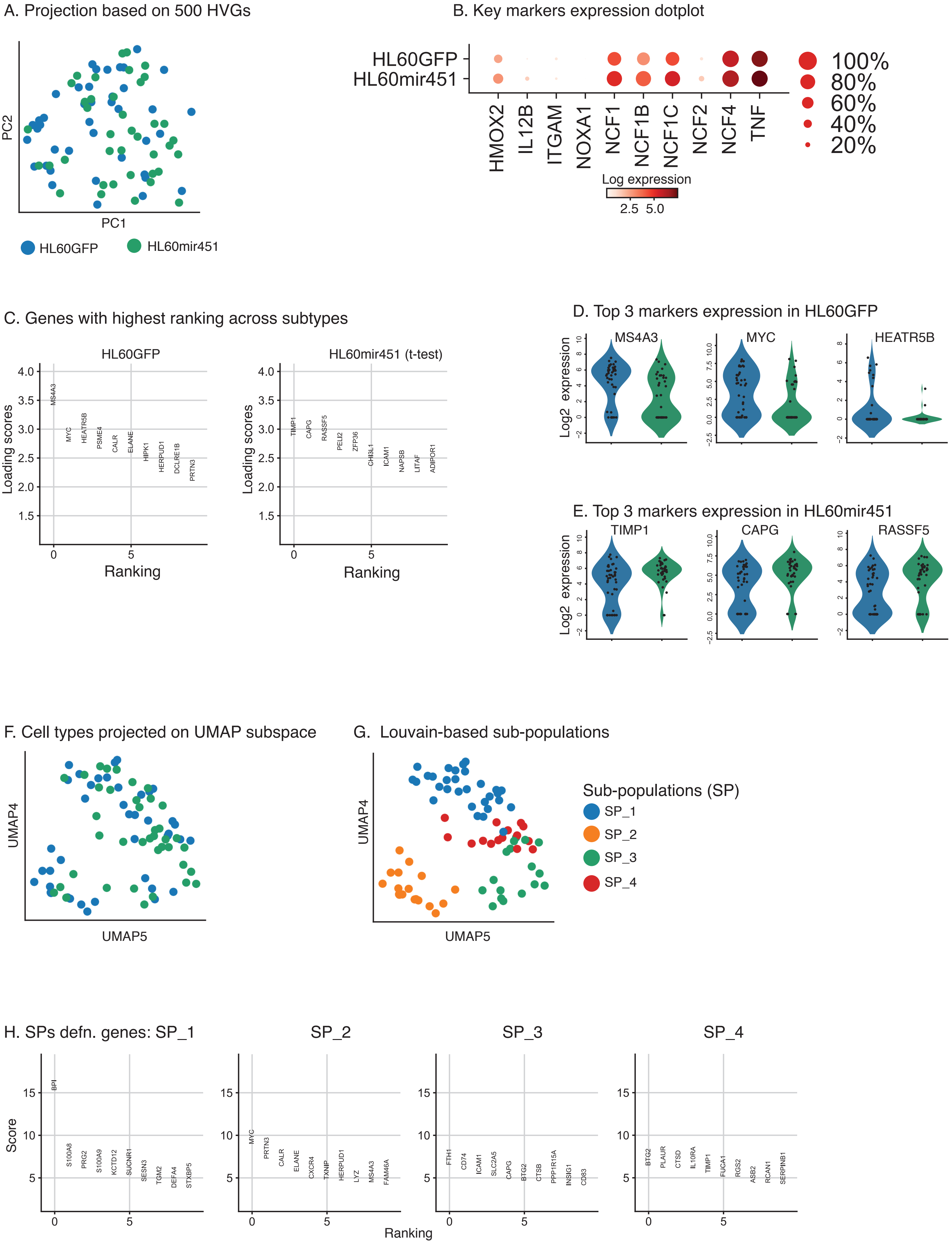

### Supplementary Figure 3: Cell types pairwise analysis (HL-60$^{scramble}$ and HL-60$^{scramble}$Bact)

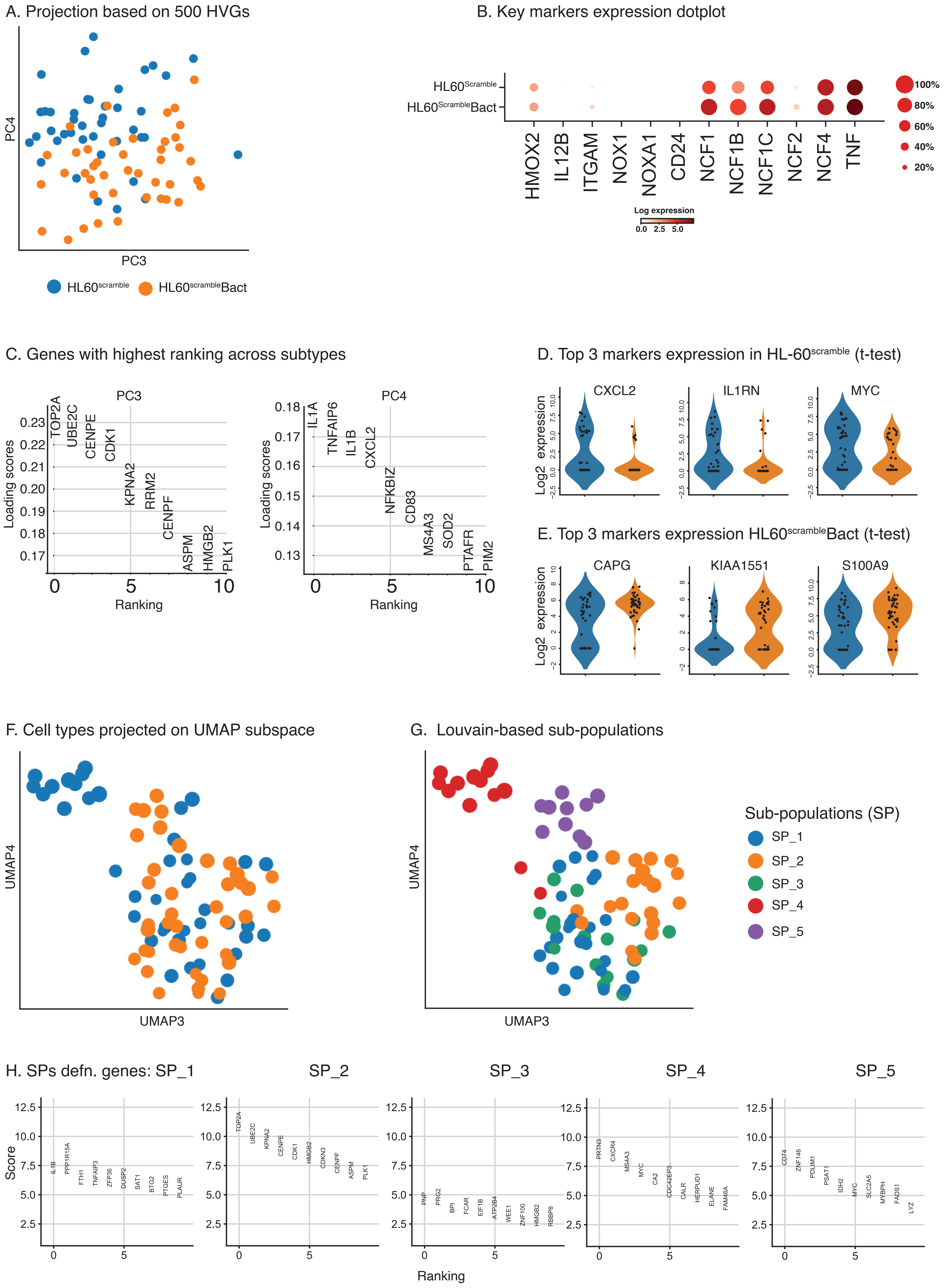

### Supplementary Fig 4: Cell types pairwise analysis ( HL-60$^{mir451a}$Bact and HL-60$^{mir451a}$)

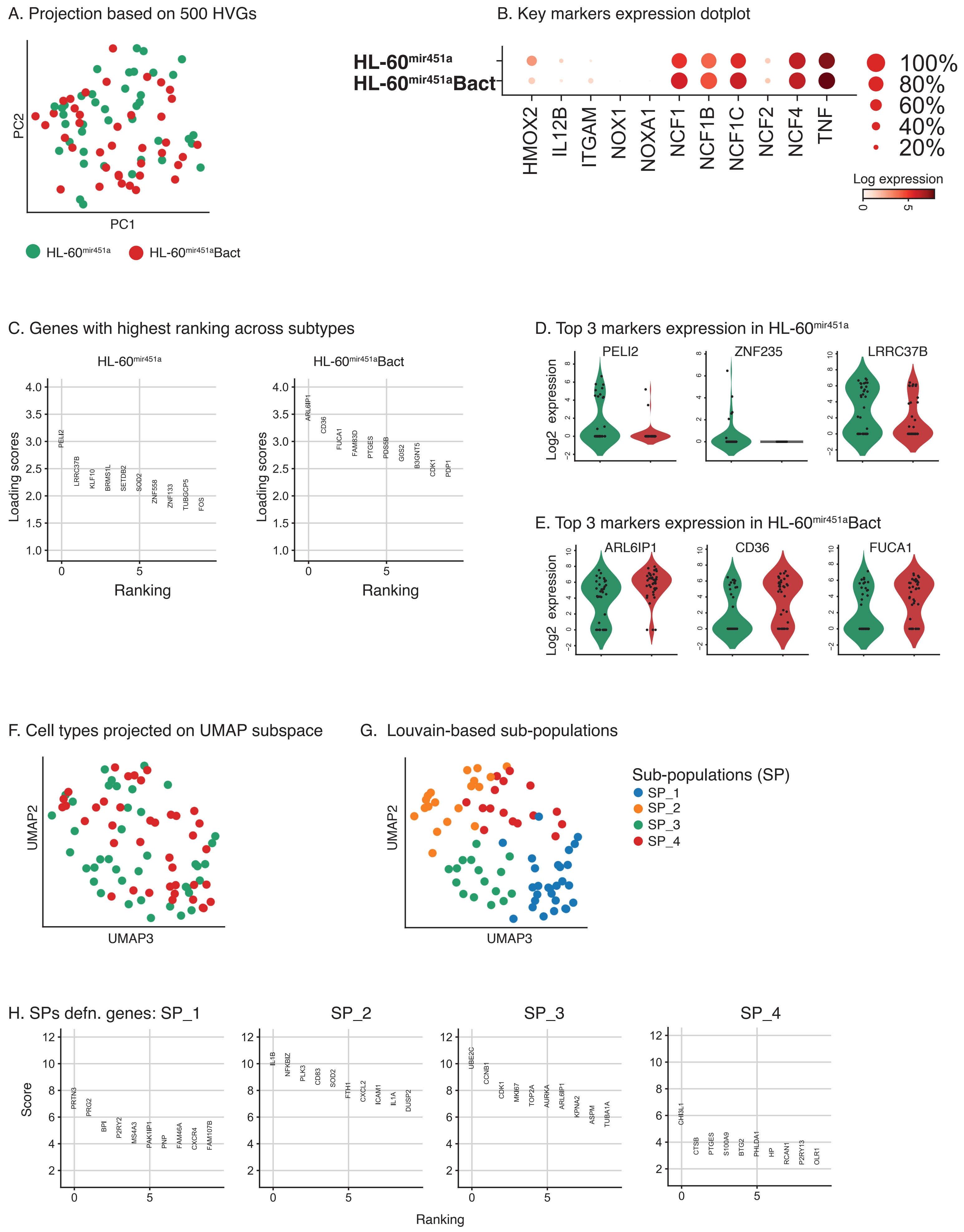

### Supplementary Fig 5: Cell types pairwise analysis (HL-60$^{Scramble}$Bact and HL-60$^{miR451a}$Bact)

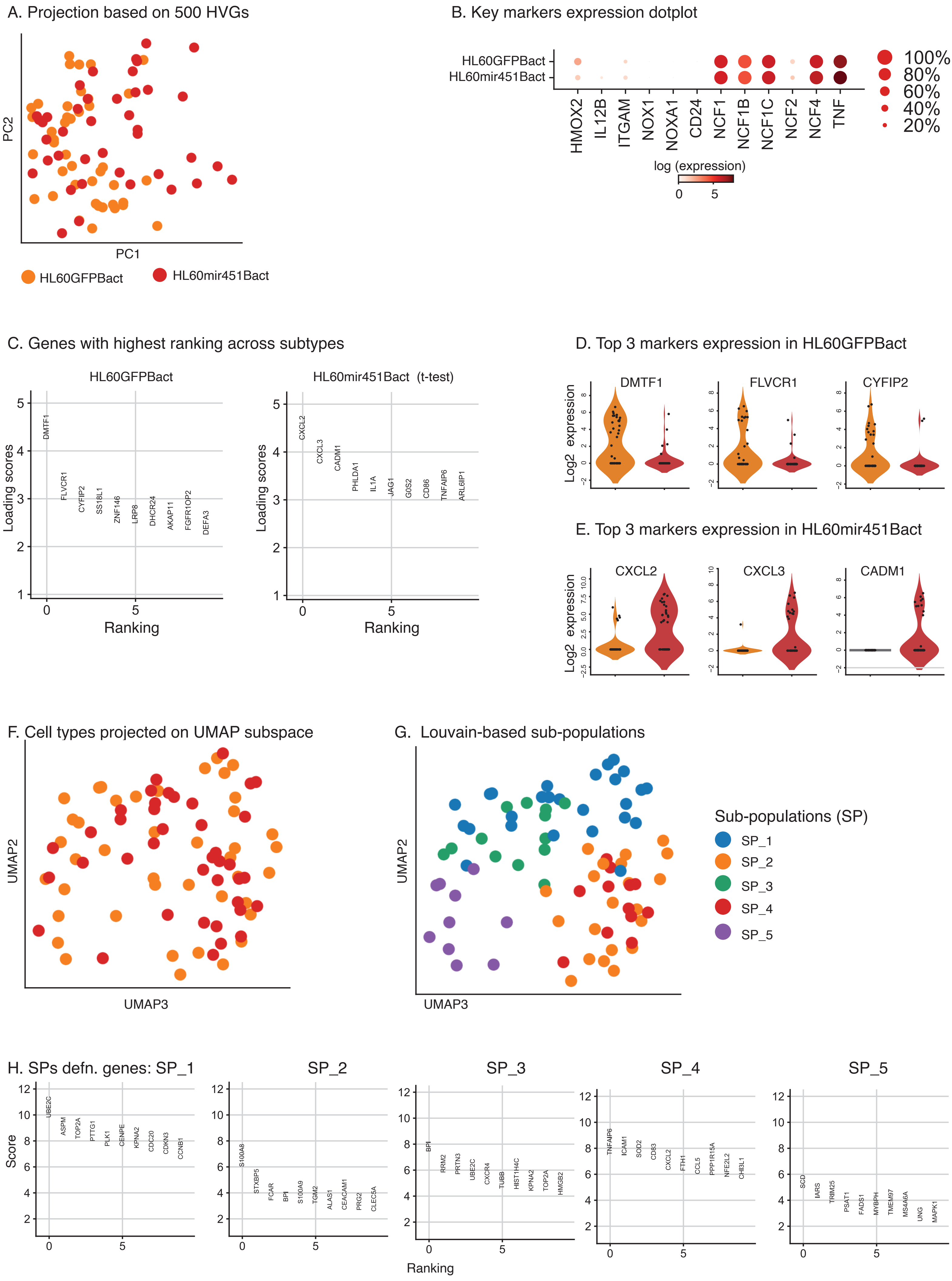

### Supplementary Fig. 6: Differential expression between HL-60$^{scramble}$Bact and HL-60$^{miR451a}$Ba

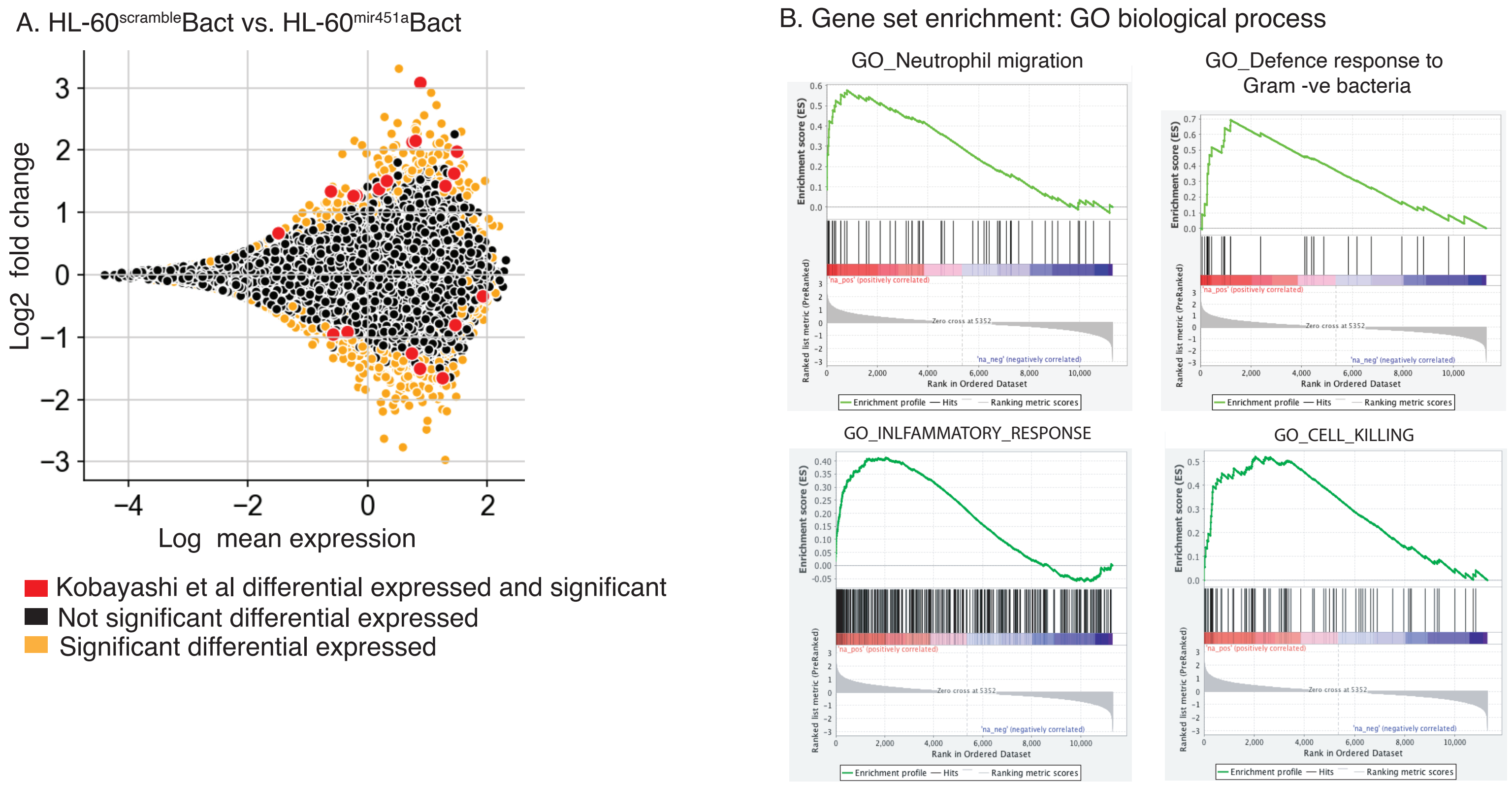

Table S1: Number of differentially expressed genes in human neutrophils after different *in-vitro* stimulations

| | **Up-regulated genes (%)** | **Down-regulated genes (%)** | **DE genes (%)** | **Unaffected genes** | **Expressed genes** |
|---|---|---|---|---|---|
| **EVs** | 3395 (25.1) | 3363 (24.8) | 6758 | 6757 | 13515 |
| **LPS** | 3832 (28.4) | 4016 (29.7) | 7848 | 5667 | 13515 |
| **LPSEV** | 3861 (28.6) | 3976 (29.4) | 7837 | 5678 | 13515 |
| **LPSEV vs Group lpsev** | 315 (2.3) | 196 (1.5) | 511 | 13004 | 13515 |

Table S2: Number of DEG in HL60miR451a after *in-vitro* stimulation with LPS at different time points

| | **LPS treatment (min)** | **Up-regulated genes (%)** | **Down-regulated genes (%)** | **DE genes (%)** | **Unaffected genes** | **Expressed genes** |
|---|---|---|---|---|---|---|
| **HL60miR451 vs HL60-GFP** | **0** | 3480 (24.4) | 3083 (21.6) | 6563 | 7713 | 14276 |
| | **90** | 39 (0.3) | 105 (0.7) | 144 | 14132 | 14276 |
| | **180** | 155 (1.8) | 261 (1.8) | 416 | 13860 | 14276 |

**SUPPLEMENTARY DATA**

**Supplementary Video S1. PMN migrating in the egg-shaped chip device**. PMN migrate towards fMLP gradient from the large egg-shaped chamber into the inner micro-chamber via the connecting channel. The inner central reservoir is about 200 µm wide while the large egg-shaped chamber is 300 µm wide. The connecting entrance channel is about 125 µm long and 10 µm wide. The circled primary neutrophils are the cells with iRBC-EVs identified by the tracking plugin on image J. The time interval between frames is 4 minutes. Scale bar is 50 µm. The nucleus of the PMN is stained with Hoechst dye (blue).

**Supplementary Video S2. *C. albicans* target growth**. Micro-patterned ***C. albicans*** (expressing a far-red fluorescent protein) spot growth in the microfluidic device assay was monitored for 10hours. The time interval between frames is 4 minutes. Scale bar is 50 µm.

**Supplementary Video S3. Untreated PMN swarming on *C. albicans* target**. PMN (blue - nucleus stained with Hoechst dye) swarm around *C. albicans* target. The *C. albicans* stained in violet dye. Isolated neutrophils are loaded on the swarming assay device at a concentration of 2.5 x $10^6$ cells/mL. The time interval between frames is 5 minutes. Scale bar is 50 µm.

**Supplementary Video S4. iRBC-EVs treated PMN swarming on *C. albicans* target**. PMN (blue - nucleus stained with Hoechst dye) swarm around *C. albicans* target. The *C. albicans* stained in violet dye. Isolated neutrophils are loaded on the swarming assay device at a concentration of 2.5 x $10^6$ cells/mL. The time interval between frames is 5 minutes. Scale bar is 50 µm.

**SUPPLEMENTARY FIGURE LEGENDS:**

**Supplementary Figure 1: Datasets overview**

A. Total reads for per cell in each treatment category (herein also referred to as cell type) $HL60^{scramble}$
B. Number of genes detected per cell grouped according to the treatment category
C. Scatter plot of total counts vs. number of genes per cell colored according to cell treatment category
D. Top 30 highly expressed genes across all cells independent of treatment
E. Top 30 highly expressed genes across all cells in $HL60^{scramble}$ group
F. Top 30 highly expressed genes across all cells in $HL60^{mir451}$ group
G. Top 30 highly expressed genes across all cells in $HL60^{scramble}$ Bact group
H. Top 30 highly expressed genes across all cells in $HL60^{mir451}$Bact group

**Supplementary Figure 2: $HL60^{scramble}$ versus $HL60^{mir451a}$**

These figures highlight the results from pairwise analysis of the two groups ($HL60^{scramble}$ and $HL60^{mir451a}$).

A. PCA projection of all $HL60^{scramble}$ and $HL60^{mir451a}$ cells based on components 1 and 2. Projection based on other components and using varying parameters failed to generate clear separation between the two cell populations
B. Dotplot of expression of some of the published key markers involved in malaria and bacterial co-infection response in the two subpopulations
C. The loading scores of top 10 genes that may be responsible for the separation of $HL60^{scramble}$ and $HL60^{mir451a}$ based on PC1 and PC2
D. The expression of top 3 genes that may be responsible for the separation of $HL60^{mir451a}$ from $HL60^{scramble}$ in PCA subspace based on the first two components 1 and 2
E. The expression of top 3 genes that may be responsible for the separation of $HL60^{mir454a}$ from $HL60^{scramble}$ in PCA subspace
F. The projection of both cell types onto UMAP based on PCA components 4 and 5
G. The 4 subpopulations (SPs) discovered based on unsupervised approach (Louvain method as implemented in Scanpy) and projected unto UMAP space
H. Some of ten genes that may be involved in defining the four SPs described above (6G)

**Supplementary Figure 3: HL60$^{Scramble}$ versus HL60$^{Scramble}$ Bact**

These figures highlight the results from pairwise analysis of the two groups (HL60$^{scramble}$ and HL60$^{scramble}$ Bact).

A. PCA projection of all HL60$^{scramble}$ and HL60$^{scramble}$Bact cells based on components 3 and 4. This was the best separation attained according to treatment of these cell types after several trials with different approaches and parameters
B. Dotplot of expression of some of the published key markers/genes involved in malaria and bacterial co-infection response
C. The loading scores of top 10 genes that may be responsible for the separation of HL60$^{Scramble}$ from HL60$^{Scramble}$Bact when projected on PC3 and PC4
D. The expression of top 3 genes that may be responsible for the separation of HL60$^{Scramble}$ from HL60$^{Scramble}$ Bact in PCA subspace based on the first two components 1 and 2
E. The expression of top 3 genes that may be responsible for the separation of HL60$^{scramble}$Bact from HL60$^{scramble}$ in PCA subspace based on the first two components 1 and 2
F. The projection of both cell types onto UMAP based on PCA components 3 and 4
G. The 5 subpopulations (SPs) discovered based on unsupervised approach (Louvain method as implemented in Scanpy) and projected unto UMAP space
H. Some of ten genes that may be involved in defining the five SPs described above (3G)

**Supplementary Figure 4: HL60$^{mir451a}$ versus HL60$^{mir451a}$Bact**

These figures highlight the results from pairwise analysis of the two groups (HL60$^{mir451a}$ and HL60$^{mir451a}$Bact).

A. PCA projection of all HL60$^{mir451a}$ and HL60$^{mir451a}$Bact cells based on components 1 and 2. Projection based on other components and using varying parameters failed to generate clear separation between the two cell populations
B. Dotplot of expression of some of the published key markers involved in malaria and bacterial co-infection response in the two subpopulations
C. The loading scores of top 10 genes that may be responsible for the separation of HL60mir451 and HL60$^{mir451a}$Bact based on PC1 and PC2
D. The expression of top 3 genes that may be responsible for the separation of HL60$^{mir451a}$ from HL60$^{mir451a}$Bact in PCA subspace based on the first two components 1 and 2.

E. The expression of top 3 genes that may be responsible for the separation of HL60$^{mir451a}$Bact from HL60$^{mir451a}$ in PCA subspace
F. The projection of both cell types onto UMAP based on PCA components 2 and 3
G. The 4 subpopulations (SPs) discovered based on unsupervised approach (Louvain method as implemented in Scanpy) and projected unto UMAP space
H. Some of ten genes that may be involved in defining the four SPs described above (8G)

**Supplementary Figure 5: HL60$^{scramble}$ Bact versus HL60$^{mir451a}$Bact**

These figures highlight the results from pairwise analysis of the two groups (HL60$^{Scramble}$ Bact and HL60$^{mir451a}$ Bact).

A. PCA projection of all HL60$^{scramble}$ Bact and HL60$^{mir451a}$ Bact cells based on components 1 and 2. Projection based on other components and using varying parameters failed to generate clear separation between the two cell populations
B. Dotplot of expression of some of the published key markers involved in malaria and bacterial co-infection response in the two subpopulations
C. The loading scores of top 10 genes that may be responsible for the separation of HL60$^{scramble}$Bact from HL60$^{mir451}$Bact based on PC1 and PC2
D. The expression of top 3 genes that may be responsible for the separation of HL60$^{scramble}$Bact from HL60$^{mir451}$Bact in PCA subspace based on the first two components 1 and 2.
E. The expression of top 3 genes that may be responsible for the separation of HL60$^{mir451}$Bact from HL60$^{scramble}$Bact in PCA subspace
F. The projection of both cell types onto UMAP based on PCA components 2 and 3
G. The 5 subpopulations (SPs) discovered based on unsupervised approach (Louvain method as implemented in Scanpy) and projected unto UMAP space
H. Some of ten genes that may be involved in defining the five SPs described above (4G)

**Supplementary Figure 6: Differential gene expression analysis between HL60$^{Scramble}$Bact and HL60$^{mir451a}$Bact**

Here are figures showing the results from the differential gene expression analysis between the two groups (HL60$^{scramble}$Bact and HL60$^{mir451a}$Bact).

A. The mean-log fold change (MA) plot of the two cell types with the significant differential expressed genes colored in orange, significant genes overlapping published ones from Kobayashi et al (Kobayashi et al. 2003; Kobayashi et al. 2002) in red and non-significant ones in black
B. Four of the significant gene ontology (GO) biological processes based on gene set enrichment analysis of log-fold change ranked genes between the two populations
